# FimX regulates type IV pilus localization via the Pil-Chp chemosensory system in *Acinetobacter baylyi*

**DOI:** 10.1101/2025.05.21.655318

**Authors:** Taylor J. Ellison, Ian Y. Yen, P. Lynne Howell, Courtney K. Ellison

## Abstract

Type IV pili (T4P) are widespread dynamic appendages required for diverse prokaryotic behaviors including twitching motility, biofilm formation, and DNA uptake leading to natural transformation. Although the components involved in T4P assembly and dynamics are largely conserved across divergent clades of bacteria, the mechanisms underlying T4P function and regulation differ significantly and remain poorly characterized outside of a select few model organisms. One understudied characteristic of T4P includes the spatial organization of T4P envelope-spanning nanomachines and how organizational patterns contribute to single cell behaviors. The bacterial species *Acinetobacter baylyi* localizes its T4P nanomachines in a unique pattern along the long axis of the cell, making it a robust model to study the mechanisms underlying the regulation of intracellular organization in single cell organisms. In this work, we find that the T4P regulatory protein FimX has been co-opted away from regulating T4P dynamics to instead control T4P positioning through a chemosensory Pil-Chp pathway. We show FimX directly interacts with the Pil-Chp histidine kinase ChpA which likely influences Pil-Chp signaling and the subsequent positioning of T4P machines. These data contribute to our understanding of how bacterial regulatory systems can evolve to function in diverse biological processes.

**SIGNIFICANCE STATEMENT:** Subcellular organization of protein complexes plays a vital role in their function in all domains of life, yet the molecular mechanisms underlying subcellular organization in bacteria remain poorly understood. This work shows the T4P regulatory component FimX directly interacts with a chemosensory signaling protein to regulate the positioning of T4P nanomachines. These data provide insight into how signaling systems can evolve to regulate diverse processes, and they establish a direct connection between environmental signaling cascades and the regulation of bacterial cell biology.

## INTRODUCTION

Many prokaryotic species encode genes to produce dynamic, filamentous appendages called type IV pili (T4P) that are required for diverse behaviors including twitching motility, DNA uptake leading to natural transformation, biofilm formation, and host cell adherence (Craig *et al*., 2019; Ellison *et al*., 2022b). Although T4P and related structures are broadly distributed across bacteria and archaea and play an important role in single-cell lifestyles, how their production and dynamics are regulated in divergent organisms remains poorly understood (Denise *et al*., 2019).

T4P dynamics are essential for their function and are driven by the activity of ATP-hydrolyzing motors (Craig *et al*., 2019; McCallum *et al*., 2019; Ellison *et al*., 2022b). Extension motors polymerize major pilin subunits that constitute the filament to drive T4P extension, while retraction motors depolymerize pilins to drive T4P retraction. In many species, accessory proteins and signaling molecules such as cyclic-3′,5′-dimeric guanosine monophosphate (cdG) play an important role in regulating T4P dynamics and function by altering motor activity (Roberge and Burrows, 2024). For example, in *Pseudomonas aeruginosa* and *Xanthomonas* spp., the small protein PilZ is required for T4P production via direct interactions with the extension motor ATPase, PilB (Alm *et al*., 1996; Guzzo *et al*., 2009). CdG also directly binds the extension motor MshE of mannose-sensitive hemagglutinin (MSHA) pili found in *Vibrio cholerae*, PilF found in *Thermus thermophilus*, and PilB in *Myxococcus xanthus* to promote motor function (Jones *et al*., 2015; Floyd *et al*., 2020; Dye *et al*., 2023; Neißner *et al*., 2025). In contrast, some proteins have been shown to directly inhibit extension motors in order to prevent T4P assembly as is the case with the inhibition of PilB by CpiA in *Acinetobacter baylyi* or the T4P-dependent phage-encoded proteins Tip, Aqs1, and Zip that inhibit T4P production in *P. aeruginosa* to prevent phage reinfection (Chung *et al*., 2014; Ellison *et al*., 2021; Shah *et al*., 2021; Taylor *et al*., 2024).

In *Xanthomonas* and *P. aeruginosa*, the cdG-binding accessory protein FimX promotes PilB extension motor activity (Guzzo *et al*., 2009; Jain *et al*., 2017; Llontop *et al*., 2021). In these species FimX contains degenerate diguanylate cyclase (GGDEF) and phosphodiesterase (EAL) domains that do not possess in vivo enzymatic activity but retain the ability to bind cdG to regulate T4P production (Guzzo *et al*., 2009; Jain *et al*., 2017). In *Xanthomonas* spp., FimX directly interacts with PilZ to promote T4P extension, while in *P. aeruginosa*, FimX instead binds to PilB. The differences between these systems highlight how accessory proteins have evolved different strategies to regulate the same processes.

To better understand the regulatory networks underlying T4P function in different species, we sought to assess the role of FimX in *A. baylyi*. Surprisingly, we found that FimX does not interact with PilB nor PilZ as it does in *P. aeruginosa* or *Xanthomonas*, respectively. In line with this observation, we show that FimX also does not promote T4P extension but instead has been co-opted to regulate T4P localization as part of the chemosensory Pil-Chp pathway that controls T4P patterning in *A. baylyi*.

## RESULTS

### FimX does not interact with PilB nor PilZ in *A. baylyi*

Studies in *P. aeruginosa* and *Xanthomonas* spp. suggested that FimX could interact with PilB or PilZ to regulate T4P extension in *A. baylyi*. To test direct interactions between FimX (ACIAD2209) and PilZ (ACIAD2360) or PilB (ACIAD0362), we performed bacterial adenylate cyclase two-hybrid (BACTH) assays. BACTH combinations were constructed by fusing the T25- or T18-fragments of the *Bordetella pertussis* adenylate cyclase to the N- or C-terminus of each protein, resulting in stable fusion constructs. Plate-based and quantitative ꞵ-galactosidase Miller assays revealed no direct interactions between FimX and PilB or PilZ, although a statistically significant, but modest, increase in ꞵ-galactosidase activity was detected between FimX and PilB compared to the empty vector control (Figure 1A, B, Supplementary Figure S1). To confirm that the null interactions were not due to protein expression levels, we performed Western blotting using an antibody against the T18 fragment and demonstrated that all three proteins were expressed (Supplementary Figure S2). We next wondered whether FimX may indirectly interact with PilB or PilZ, which is more easily detected by *in vivo* coimmunoprecipitation assays that can detect interactions between components within multiprotein complexes. We thus created strains encoding protein fusions *fimX-GFP* and *3xFLAG-pilB* or *3xFLAG-pilZ* for coimmunoprecipitation assays in *A. baylyi*. The *3xFLAG-pilB* allele was previously found to be fully functional by natural transformation assays (Ellison *et al*., 2021), and the *3xFLAG-pilZ* strain was likewise able to undergo natural transformation at wildtype levels (Supplemental Figure S3). Coimmunoprecipitation assays using ⍺-FLAG magnetic beads and 3xFLAG-PilZ or 3xFLAG-PilB as bait resulted in no FimX-GFP signal in the elution fraction, suggesting FimX does not indirectly interact with either PilZ or PilB (Figure 1C).

**Figure 1.**
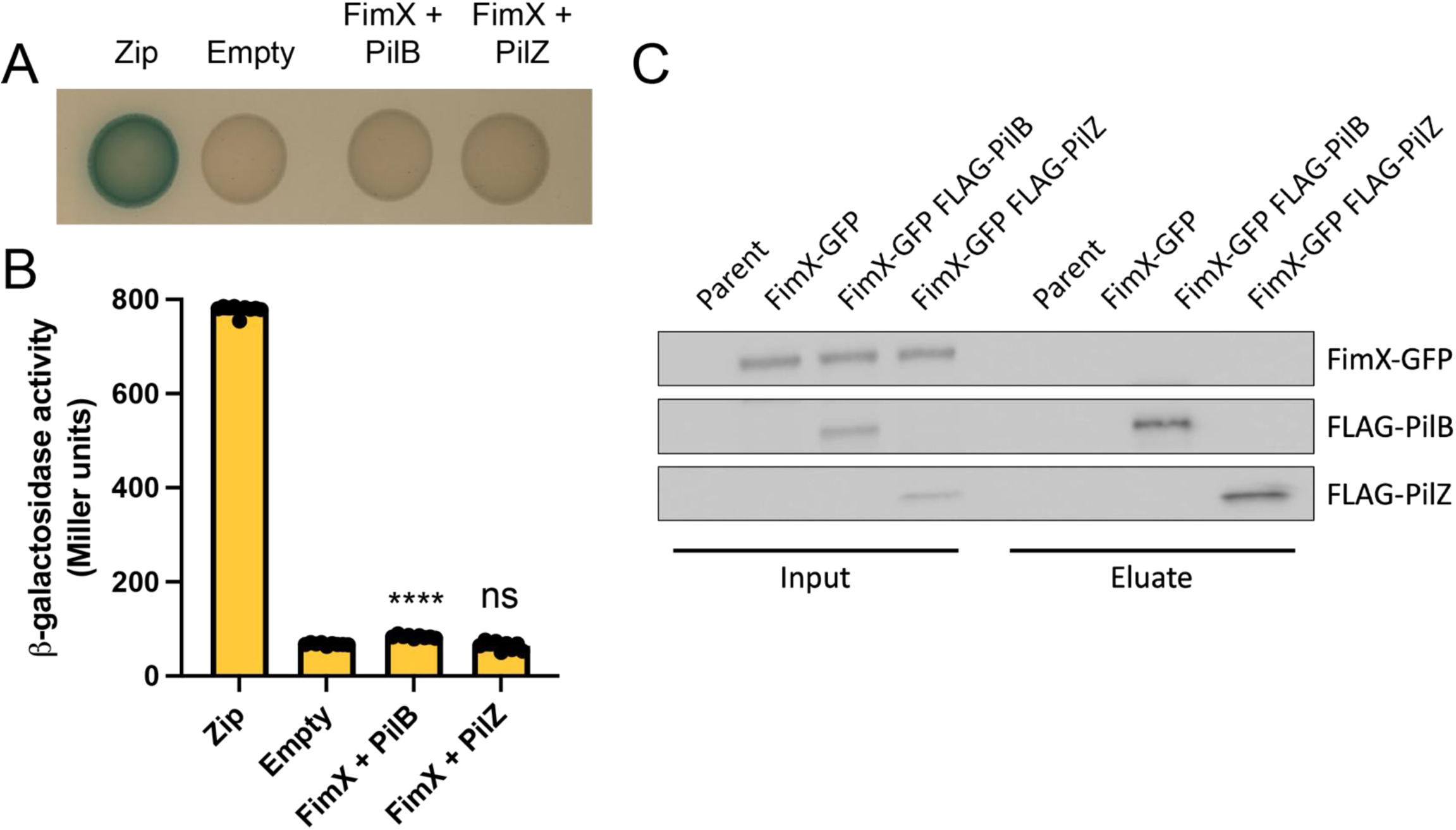
FimX does not interact with PilB nor PilZ in *A. baylyi.* A) BACTH interactions between FimX and PilB/PilZ fused N-terminally to the T25 and T18 domains of adenylate cyclase, respectively, following incubation at 30 °C for 48 hours and subsequently at 4 °C for 24 hours on LB agar plates containing X-Gal. Leucine zipper motif (zip) fused to T18 and T25 was used as positive control while empty vectors of T18 and T25 alone (empty) was the negative control. B) Quantification of ꞵ-galactosidase activity in Miller units for the same interactions depicted in A. The data show three biological replicates each with three technical replicates. Error bars show mean ± SD and statistical significance was determined using Dunnett’s multiple comparisons test against the empty vector control. \*\*\*\**p* < 0.0001. ns, not significant. C) Western blot showing coimmunoprecipitation experiments where 3xFLAG-PilB or 3xFLAG-PilZ was used as the bait protein to test for interaction with FimX-GFP as the prey protein. Blot was probed using both ⍺-FLAG and ⍺-GFP antibodies to detect both proteins simultaneously.

### Deletion mutants lacking *fimX* make mislocalized T4P

Because FimX does not appear to regulate T4P extension via PilB or PilZ, we wondered if FimX may regulate T4P assembly via an alternative mechanism. T4P and their dynamics are required for DNA uptake leading to natural transformation, and T4P mutants defective in T4P synthesis or retraction are unable to undergo efficient natural transformation (Seitz and Blokesch, 2013; Ellison *et al*., 2018). We thus performed natural transformation assays to assess the role of FimX in T4P production. As expected, a major pilin deletion strain Δ*comP* fell below the limit of detection (Figure 2A). However, the parent strain and the Δ*fimX* mutant exhibited similar transformation frequencies, suggesting that FimX does not influence T4P production (Figure 2A).

**Figure 2.**
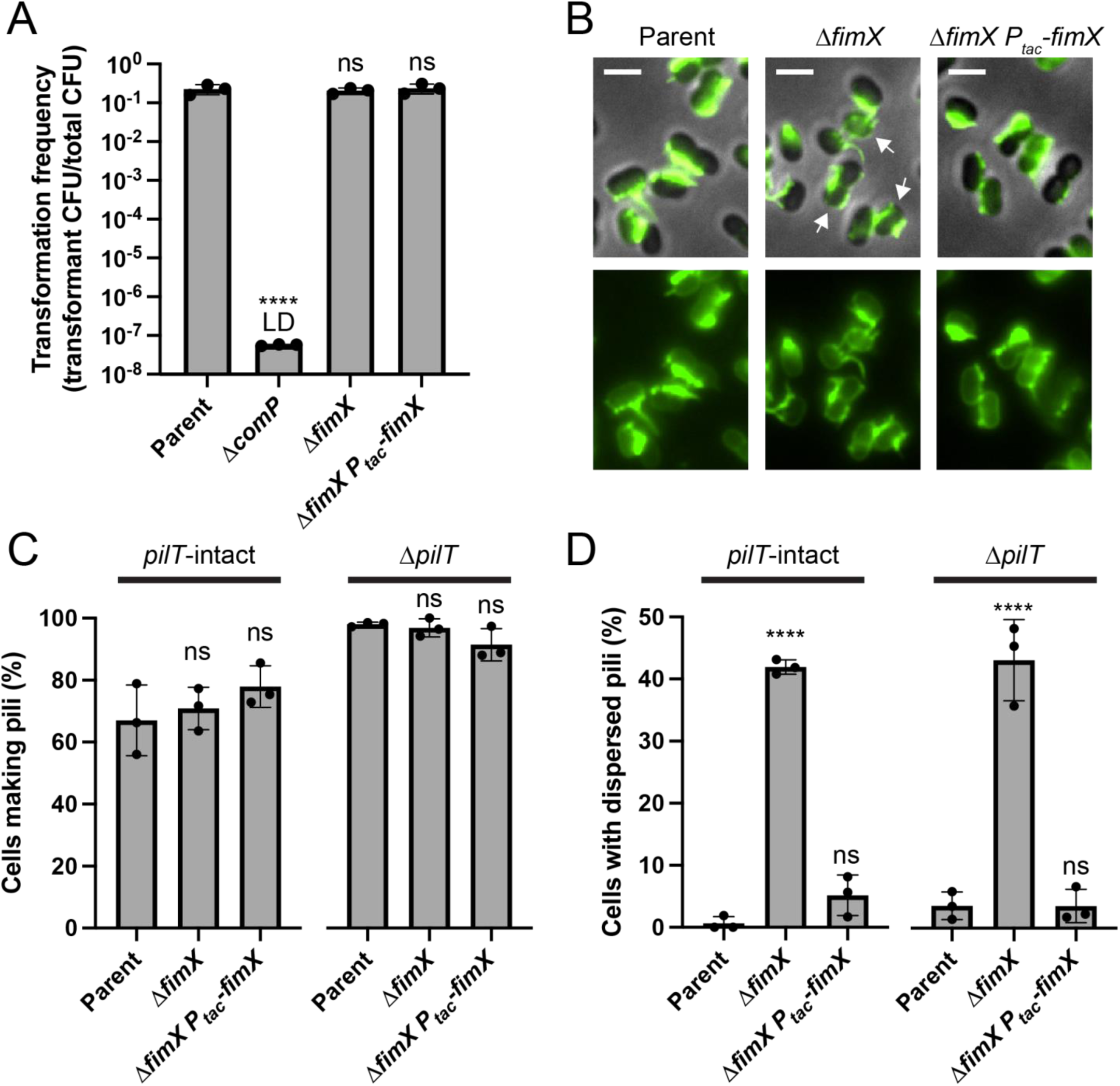
Deletion mutants of *fimX* make mislocalized T4P. A) Natural transformation assays of indicated strains. LD, limit of detection. Statistical significance was determined using log-transformed transformation frequencies. B) Representative microscopy images of indicated strains in a Δ*pilT* retraction motor deletion parent strain labeled with AF488-mal. Arrows indicate cells with mislocalized T4P. Scale bars, 2 µm. C) Percent of piliated cells in indicated strains. D) Percent of cells with mislocalized or dispersed T4P in indicated strains. Cells with dispersed T4P were defined as cells that had T4P on both long-axis sides of the cell body. For panels A,C,D, each data point represents a biological replicate (n = 3), and bar graphs indicate the mean ± SD. A minimum of 50 cells were analyzed per replicate. Statistical significance was determined using Dunnett’s multiple comparisons test against the parent strain for each data set. \*\*\*\**p* < 0.0001. ns, not significant.

Because FimX does not promote T4P synthesis, we wondered whether it may be involved in regulating another aspect of T4P function in *A. baylyi*. Previous work highlighted that wildtype *A. baylyi* produces T4P in a line along the long axis of the cell, and deletion mutants in the chemosensory Pil-Chp pathway produce T4P that are dispersed across the cell body (Ellison *et al*., 2022a). To test whether FimX may play a role in T4P localization, we used fluorescence microscopy to image T4P in single cells. T4P can be fluorescently labeled via the introduction of a cysteine residue into the major pilin subunit for subsequent labeling with thiol-reactive maleimide dyes (Ellison *et al*., 2017, 2019, 2021). T4P were produced at similar levels to the parent strain (Figure 2B, C), but Δ*fimX* mutants exhibited mislocalized or dispersed T4P (Figure 2B, D). To more easily assess T4P phenotypes, we also analyzed Δ*pilT* mutants lacking the retraction motor which results in retraction-deficient mutants and consequent hyperpiliation. Δ*pilT* strains exhibited similar phenotypes to their *pilT*-intact counterparts, confirming that FimX does not promote T4P assembly and is instead involved in regulating T4P localization (Figure 2C, D). The mislocalization of T4P in Δ*fimX* mutants suggests FimX participates in Pil-Chp signaling (Ellison *et al*., 2022a).

### FimX is part of the Pil-Chp chemosensory system

The Pil-Chp pathway shares homology to flagellar chemotaxis systems (Figure 3A) and is best-characterized in *P. aeruginosa* where it regulates T4P synthesis through T4P-mediated surface-sensing (Siryaporn *et al*., 2014; Luo *et al*., 2015; Persat *et al*., 2015; Webster *et al*., 2022). Although the Pil-Chp pathway in *P. aeruginosa* possesses a methyl-accepting chemotaxis homologue PilJ that is important for its function, PilJ is not required for T4P linear localization in *A. baylyi* unlike most of the other Pil-Chp components (Ellison *et al*., 2022a). In *P. aeruginosa*, signaling from PilJ is thought to initiate phosphotransfer from the histidine kinase ChpA (homologous to CheA in flagellar chemotaxis systems) to the response regulator PilG (a CheY homologue), followed by further downstream signaling to induce T4P extension in response to mechanical contact (Silversmith *et al*., 2016; Kühn *et al*., 2021). In *P. aeruginosa*, PilG is connected to the T4P machinery-affiliated component FimV through the scaffolding protein FimL to regulate T4P production (Inclan *et al*., 2016). However, in *A. baylyi* these components similarly interact to regulate T4P machinery localization instead of T4P extension (Ellison *et al*., 2022a). Consistent with the Pil-Chp system playing a role in T4P machinery localization, previous work also showed that PilG, FimL, and FimV localize to sites of T4P synthesis in *A. baylyi* (Ellison *et al*., 2022a). We thus examined the localization of FimX-mRuby3 in relation to sites of T4P production. Like all other FimX C-terminal fusion strains used throughout this work, FimX-mRuby3 was fully functional with only ∼5% of cells producing dispersed T4P compared to ∼55% in the Δ*fimX* control (Supplemental Figure S4). Microscopy analysis revealed that FimX-mRuby3 localizes to the base of T4P similar to other Pil-Chp components, and approximately 90% of cells with T4P exhibited colocalization between FimX-mRuby3 and labeled T4P (Figure 3B).

**Figure 3.**
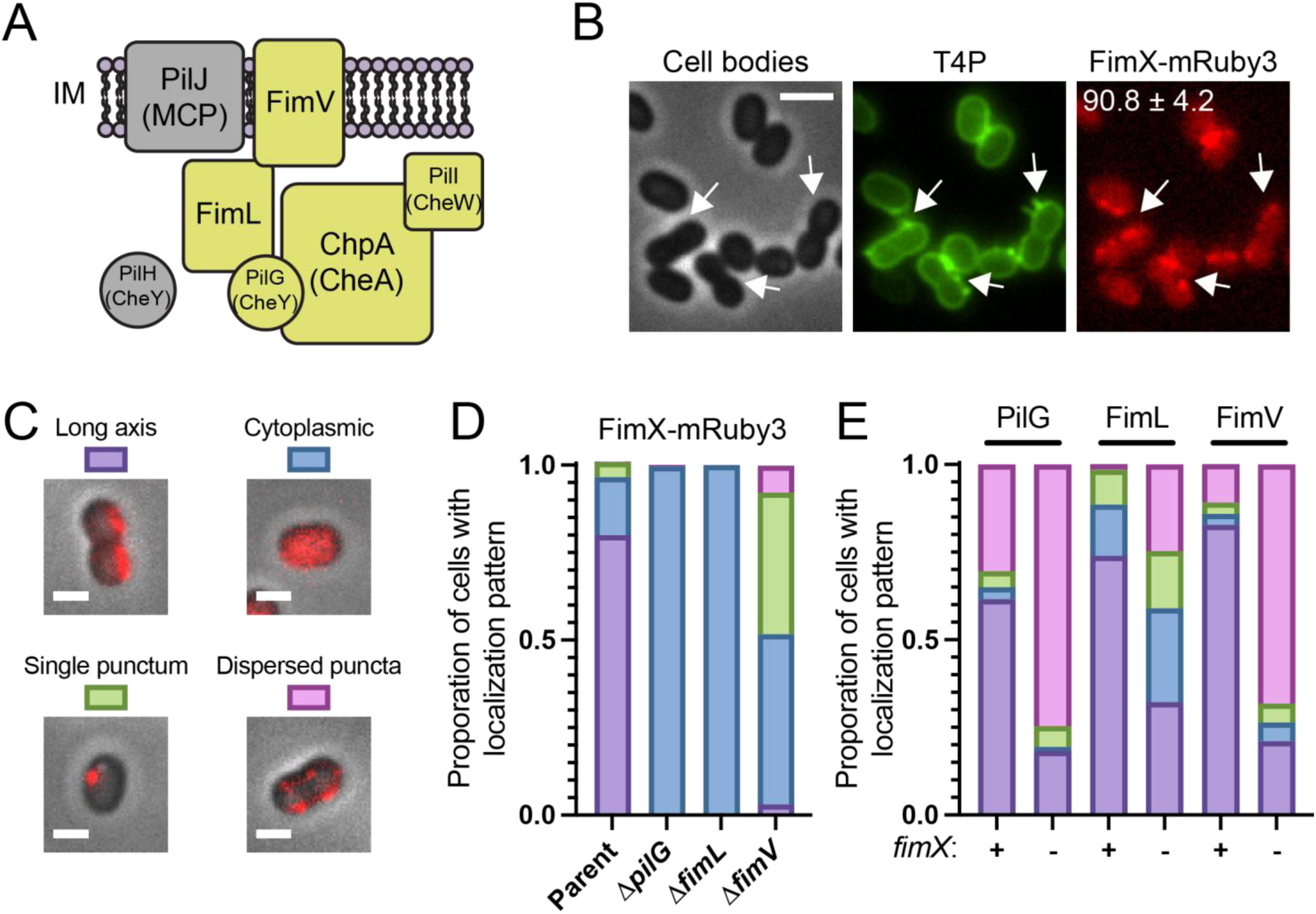
FimX is part of the Pil-Chp chemosensory system. A) Schematic of the Pil-Chp components found in *A. baylyi* with analogous flagellar chemotaxis protein names in parentheses. Deletions of components colored gold cause dispersed T4P localization while deletion of components in gray have no effect on localization of T4P. IM, inner membrane. B) Representative microscopy images of strain containing FimX-mRuby3 with AF488-mal labeled. Scale bar, 2 µm. Arrows indicate FimX-mRuby3 localizing to the base of T4P. Number in top right corner indicates the average percent of piliated cells that have FimX-T4P colocalization. C) Representative microscopy images of single cells with localization patterns quantified in D and E. Scale bars, 1 µm. D) Quantification of FimX-mRuby3 localization in indicated strains. E) Quantification of localization patterns of PilG-mRuby3, FimL-mCherry, or FimV-mRuby3 in the presence or absence of *fimX*. Data for D and E are shown as the average proportion of each localization pattern shown in C from three biological replicates. A minimum of 50 total cells were assessed for each biological replicate.

One complicating feature of the Pil-Chp system is the interdependent feedback signaling between components, and the mechanism underlying this interdependency remains unclear. PilG and FimL require each other to associate with FimV and both are required for FimV to linearly localize along the long axis and vice versa, demonstrating that complex signaling between these components is essential to establish linear T4P patterning (Ellison *et al*., 2022a). To test whether FimX may also be regulated by signaling via Pil-Chp components, we assessed FimX localization in Δ*pilG*, Δ*fimL*, or Δ*fimV* mutants. As done previously, we classified localization patterns of fluorescently tagged proteins into four categories: 1) long-axis localization, 2) cytoplasmically diffuse signal, 3) fluorescence localized into a single punctum, or 4) multiple, fluorescent puncta localized to opposite sides of the cell body (Figure 3C). In all mutants, FimX no longer localized to the long axis of the cell, and instead generally exhibited cytoplasmic localization (Figure 3C, D). The mislocalization of FimX also suggested that FimX may be involved in signaling to regulate the localization of other Pil-Chp components. Deletion of *fimX* reduced the proportion of cells with linearly positioned PilG, FimL, and FimV, supporting the model that FimX modulates the Pil-Chp signal cascade (Figure 3C, E).

### *A. baylyi* FimX functions independently of cdG and GTP

A critical aspect of FimX in regulating T4P function in both *P. aeruginosa* and *Xanthomonas* spp. is its ability to bind cdG via its EAL domain. A Foldseek analysis coupled with AlphaFold3 prediction suggested FimX from *A. baylyi* (*Ab*FimX) exhibits similar domain organization to the *P. aeruginosa* homologue (*Pa*FimX), containing degenerate CheY-like receiver (REC) and Per-ARNT-Sim (PAS) domains at the N-terminus followed by a degenerate GGDEF-like and EAL domains at the C-terminus (Figure 4A, Supplemental Figure S5) (Gough *et al*., 2001; Pandurangan *et al*., 2019; Abramson *et al*., 2024; van Kempen *et al*., 2024). *Pa*FimX and *Ab*FimX share 21% overall sequence identity and 39% sequence similarity. In *Xanthomonas*, cdG binding is dependent on canonical “EAL” residues within the EAL domain, while in *P. aeruginosa* these residues have diverged slightly to “EVL” but retain cdG binding with high affinity (Navarro *et al*., 2009). Sequence alignment between *Pa*FimX and *Ab*FimX revealed further divergence of these residues to “EVT” in *A. baylyi*, suggesting that this FimX homolog may be incapable of binding cdG (Figure 4A, Supplemental Figure S6). To test whether *Ab*FimX could bind cdG, we recombinantly purified FimX from both species and assessed cdG binding using isothermal titration calorimetry (ITC). While *Pa*FimX bound cdG with nanomolar affinity (dissociation constant K_D_ = 144 nM) consistent with previously reported data (Navarro *et al*., 2009), no interaction between *Ab*FimX and cdG could be detected (Figure 4B). Superimposition of the crystal structure of cdG bound to the EAL domain of *Pa*FimX with the Alphafold3 predicted EAL domain of *Ab*FimX highlighted drastic alterations in residues that mediate the nucleotide base including Y673, E654, F652, and H533 in *Pa*FimX that diverged to S682, N663, E661, and N540 in *Ab*FimX, respectively (root mean square deviation of 1.2 Å over 147 Cα atoms) (Figure 4C, Supplemental Figure S6) (Navarro *et al*., 2009). Notably, L477 of the “EVL” motif and R479 in *Pa*FimX that support the phosphate ring have changed to T485 and G487, respectively, in *Ab*FimX, abrogating its ability to mediate the negatively charged phosphates (Figure 4C, Supplemental Figure S6).

**Figure 4.**
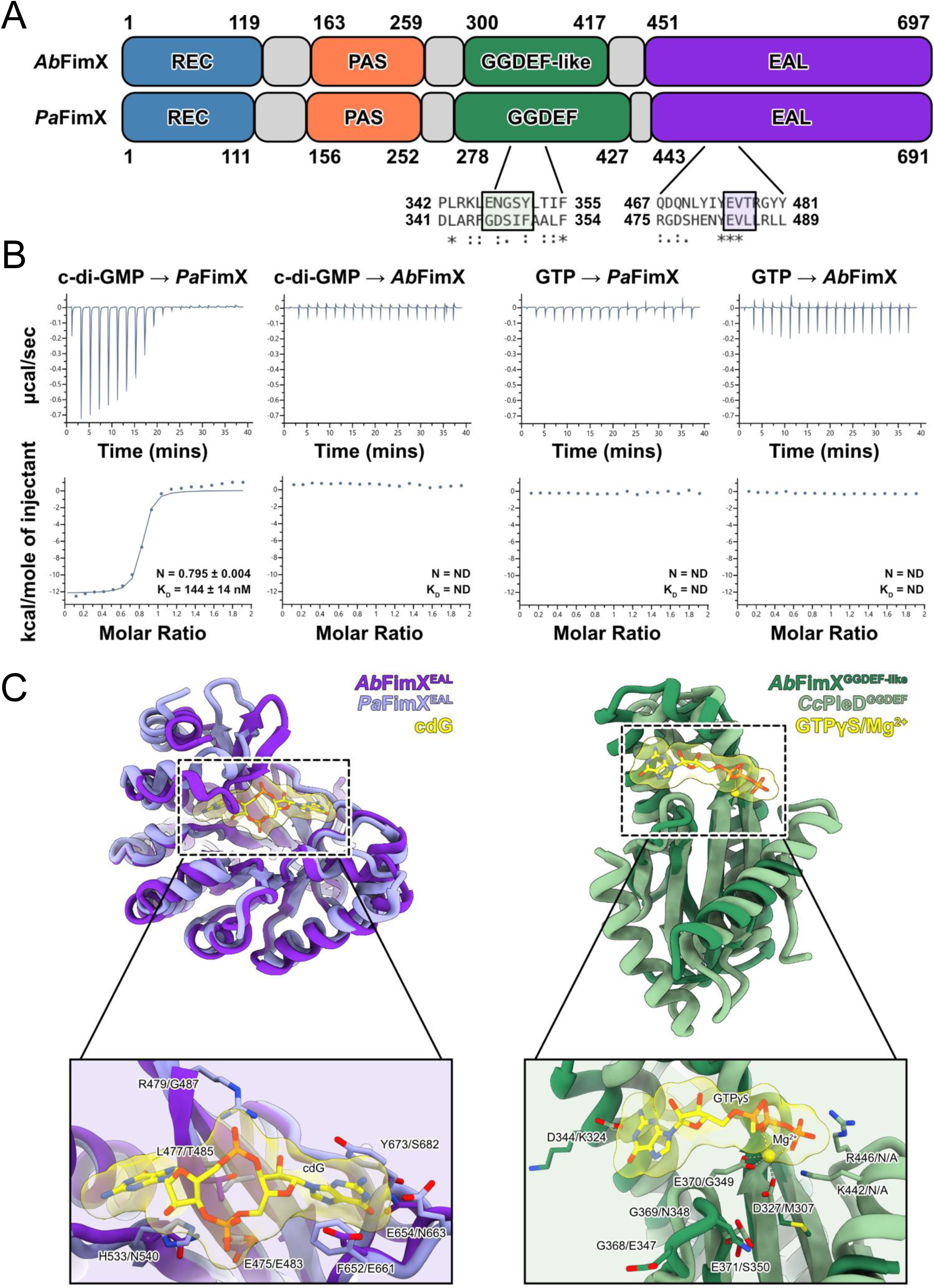
*A. baylyi* FimX does not bind cdG nor GTP. A) Schematic of *A. baylyi* (*Ab*FimX) and *P. aeruginosa* (*Pa*FimX) FimX domain organization from Foldseek and AlphaFold3 modeling. In *Ab*FimX, the blue region encompasses a receiver domain (REC); orange region encompasses a Per-ARNT-Sim (PAS) domain; green region encompasses a GGDEF-like domain; purple region encompasses a phosphodiesterase domain (EAL). Black-boxed region shows sequence alignment of the degenerate GGDEF and EAL motifs between *Ab*FimX and *Pa*FimX. B) Calorimetric titrations of 250 μM cdG or 1 mM GTP into 25 μM or 100 μM *Pa* and *Ab* FimX, respectively. Top panel depicts the heat release from injections while the bottom panel shows the normalized fitted binding curve. The stoichiometry (N) and the dissociation constant (KD) are displayed if binding was observed. ND, not determined. C) (left) Superimposition of the crystal structure of the cdG-bound EAL domain from *Pa*FimX (light purple) (PDB: 3HV8) onto the AlphaFold3 predicted EAL domain of *Ab*FimX (dark purple). Bottom panel depicts the close-up view of the active site with residues in *Pa*FimX involved in cdG binding shown on the left and equivalent residues that diverged in *Ab*FimX highlighted on the right. (right) Superimposition of the crystal structure of the GTPγS-bound GGDEF domain from *Caulobacter crescentus* PleD (*Cc*PleD) (light green) (PDB: 2V0N) onto the AlphaFold3 predicted GGDEF-like domain of *Ab*FimX (dark green). Bottom panel depicts the close-up view of the nucleotide binding pocket with residues in *Cc*PleD involved in GTPγS/Mg^2+^ binding shown on the left and equivalent residues that diverged in *Ab*FimX depicted on the right.

Alphafold3 predicted a GGDEF-like domain in *Ab*FimX with three rather than the typical five-stranded ꞵ-sheet fold, along with divergence of the “GGDEF” sequence motif to “ENGSY” upon alignment with *Pa*FimX (Figure 4A, Supplemental Figures S5, S6). Given the role of GGDEF-containing proteins in synthesizing cdG from guanosine triphosphate (GTP), we additionally probed for GTP binding to *Pa*FimX and *Ab*FimX using ITC and found that neither protein bound GTP even at high protein concentrations (Figure 4B). Superimposition of the crystal structure of GTPγS bound to the GGDEF domain of the *Caulobacter crescentus* response regulator PleD (*Cc*PleD) with the AlphaFold3 predicted GGDEF-like domain from *Ab*FimX revealed similarly nonconservative changes in residues that coordinate the magnesium ion and GTPγS including D344, D327, and E370 of the “GGEEF” motif of *Cc*PleD to K324, M307, and G349 of *Ab*FimX, respectively (root mean square deviation of 5.2 Å over 44 Cα atoms) (Figure 4C) (Wassmann *et al*., 2007). In addition, the α-helix that houses R446 and K442 of *Cc*PleD to mediate the γ-phosphate is truncated in *Ab*FimX and thereby missing these positively charged residues (Figure 4C). Navarro and colleagues reported similar residue alterations in *Pa*FimX that coordinate the magnesium ion and γ-phosphate, which explains the lack of GTP binding (Figure 4B) (Navarro *et al*., 2009). Together, our biophysical and structural analyses suggest that the lack of binding to cdG in *Ab*FimX further supports its functional divergence towards T4P localization.

### cAMP signaling does not influence T4P production nor localization

Because of its role in Pil-Chp signaling, it is possible that other Pil-Chp signaling molecules may play an important role in FimX function. In *P. aeruginosa*, Pil-Chp signaling is linked to increasing cyclic adenosine monophosphate (cAMP) levels via the adenylate cyclase CyaB to upregulate hundreds of genes (including those involved in T4P production) through the transcriptional regulator Vfr (Fulcher *et al*., 2010; Luo *et al*., 2015). We thus wondered whether cAMP levels or Vfr activity may regulate T4P localization via FimX in *A. baylyi*. Deletion of the genes encoding either CyaB (ACIAD1397) or Vfr (ACIAD1262) in *A. baylyi* resulted in no alteration in T4P localization (Figure 5A, B). Assessment of piliation similarly demonstrated no change in T4P produced in Δ*cyaB* or Δ*vfr* mutants (Figure 5A, C). Together, these results demonstrate that cAMP levels and Vfr-regulated genes likely do not regulate FimX function or T4P production in *A. baylyi*.

**Figure 5.**
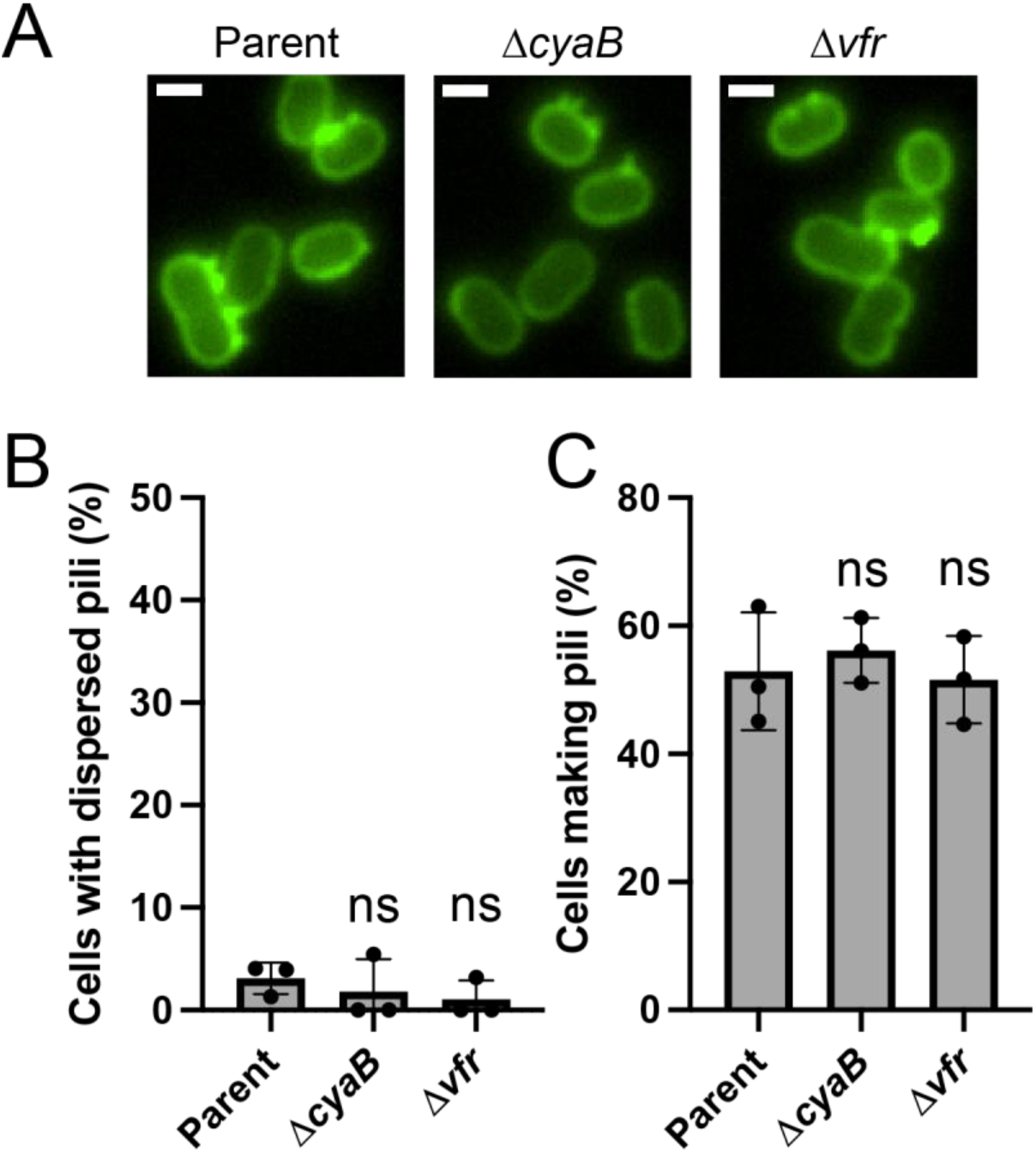
cAMP signaling factors do not influence T4P production nor localization. A) Representative microscopy images of indicated strains labeled with AF488-mal with background fluorescence subtracted. Scale bars, 1 µm. B) Percent of cells with mislocalized or dispersed T4P in indicated strains. C) Percent of piliated cells in indicated strains. Each data point represents a biological replicate (n = 3), and bar graphs indicate the mean ± SD. A minimum of 50 cells were analyzed per replicate. Statistical significance was determined using Dunnett’s multiple comparisons test against the parent strain for each data set. ns, not significant.

### FimX interacts with ChpA to regulate linear T4P localization via the Pil-Chp pathway

To identify interaction partners that could provide insight into FimX function in Pil-Chp signaling, we performed a coimmunoprecipitation experiment using a functional FimX-3xFLAG strain (Supplemental Figure S4) followed by mass spectrometry analysis as done previously (Ellison *et al*., 2022a). Mass spectrometry revealed a potential interaction between FimX and ChpA (Supplementary Data File 1), suggesting that FimX may influence Pil-Chp signaling via the ChpA sensor kinase. In support of this hypothesis, coimmunoprecipitation experiments performed using ChpA-3xFLAG as the bait protein pulled down FimX (Figure 6A). To determine whether ChpA and FimX directly interact, we next performed BACTH assays using plated-based and quantitative ꞵ-galactosidase Miller approaches (Figure 6B, C, Supplemental Figure S7). ChpA and FimX were able to directly interact in the absence of other Pil-Chp components. ChpA and PilG both regulate T4P localization in *A. baylyi*, and phosphotransfer between ChpA and PilG is an essential component of Pil-Chp signaling (Silversmith *et al*., 2016). These results suggest that FimX participates in the Pil-Chp pathway via direct interaction with ChpA to modulate signaling required for linear T4P localization.

**Figure 6.**
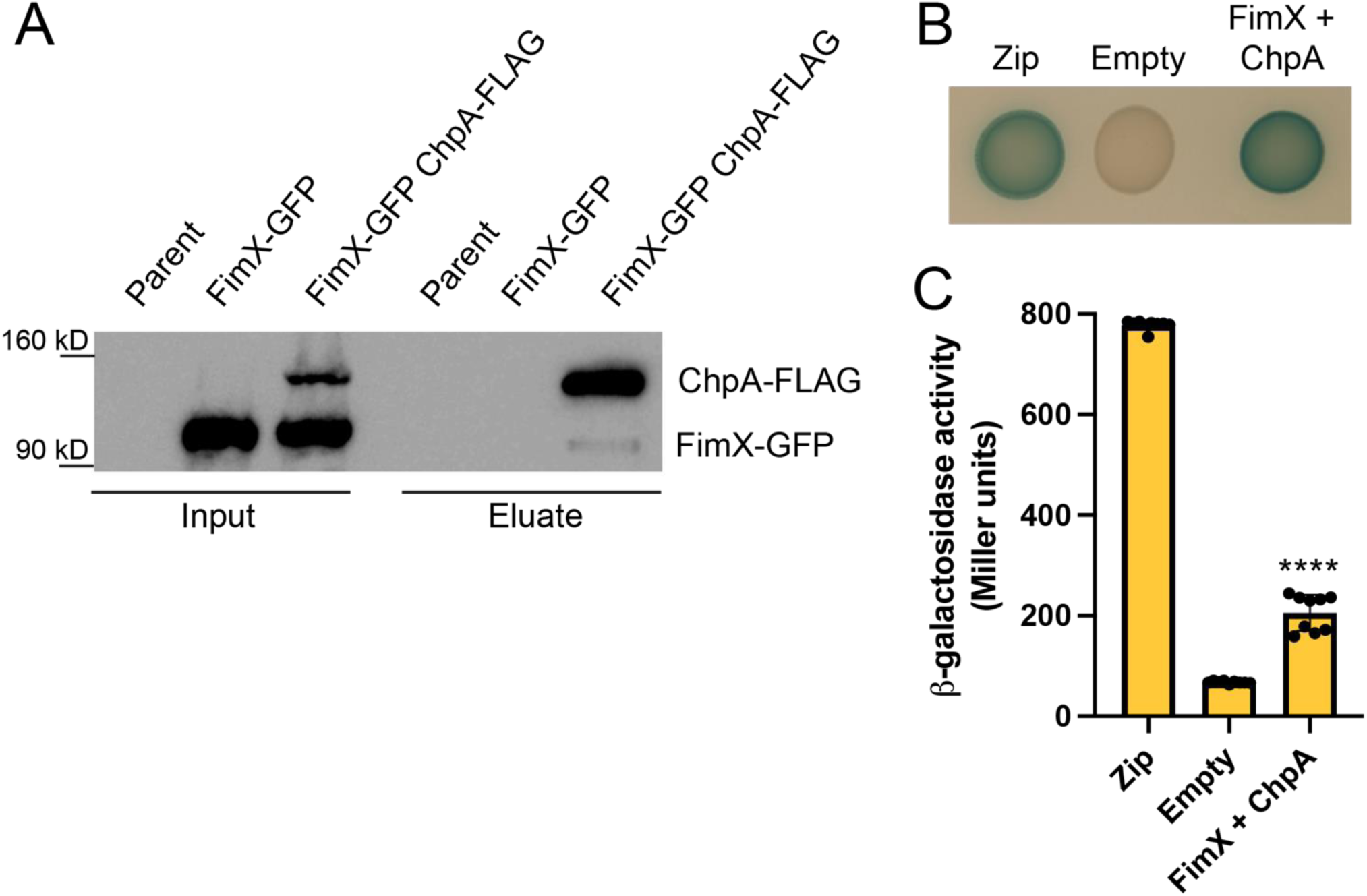
FimX directly interacts with ChpA. A) Western blot showing coimmunoprecipitation experiments where ChpA-FLAG was used as the bait protein to test for interaction with FimX-GFP as the prey protein. The blot was probed using both ⍺-FLAG and ⍺-GFP antibodies to detect both proteins simultaneously. B) BACTH interaction between FimX and ChpA fused N-terminally to the T25 and T18 domains of adenylate cyclase, respectively, following incubation at 30 °C for 48 hours and subsequently at 4 °C for 24 hours on LB agar plates containing X-Gal. Leucine zipper motif (zip) fused to T18 and T25 was used as positive control while empty vectors of T18 and T25 alone (empty) was the negative control. C) Quantification of ꞵ-galactosidase activity in Miller units for the same interactions depicted in B. The data show three biological replicates each with three technical replicates. Error bars show mean ± SD and statistical significance was determined using Dunnett’s multiple comparisons test against the empty vector control. \*\*\*\**p* < 0.0001.

## DISCUSSION

In this manuscript, we demonstrate that the accessory protein FimX has been co-opted to regulate T4P localization. This is in contrast to *P. aeruginosa* where binding of FimX to PilB promotes T4P extension dependent on cdG (Jain *et al*., 2017). Our protein interaction studies in *A. baylyi* demonstrated no binding between PilB and FimX, presumably due to its inability to bind cdG at the EAL domain due to residue divergence (Figures 1, 4). This could explain the functional divergence of FimX away from promoting T4P synthesis, consistent with the observation that deletion strains of *A. baylyi fimX* retain piliation and natural transformation capabilities (Figure 2). The inability of *A. baylyi* FimX to bind GTP, further suggests a shift away from cdG usage (Figure 4). Our finding that FimX directly interacts with ChpA and affects the localization of other Pil-Chp proteins indicates participation in coordinating the Pil-Chp signaling cascade (Figures 3, 6).

Interestingly, FimX homologues contain a conserved REC domain that shares homology to CheY, but this domain lacks the phosphorylated aspartate that is essential for signaling. However, it retains the two aspartate residues at the N-terminus (In *A. baylyi* FimXD21 analogous to CheYD12 and FimXD22 analogous to CheYD13) that bind metal ions essential for coordinating phosphorylation (Bellsolell *et al*., 1994). It is thus possible that the REC domain of FimX may assist in coordinating phosphotransfer between Pil-Chp components or within the many phosphotransfer domains found in ChpA (Silversmith *et al*., 2016). Divergence of ChpA in *A. baylyi* may also explain its ability to interact with FimX. *P. aeruginosa* ChpA is a large ∼270 kD protein compared to the relatively smaller ∼160 kD ChpA in *A. baylyi*. These changes to ChpA may eliminate or enable new protein interactions as is the case for FimX and ChpA reported here. Although our data show that FimX in *A. baylyi* does not bind cdG, mass spectrometry analysis of FimX-3xFLAG coimmunoprecipitation eluates identified a diguanylate cyclase GGDEF domain-containing protein (ACIAD2242) as a potential FimX interaction partner. ACIAD2242 is transcribed opposite of the secondary extension motor, TfpB, thus an alternative explanation is that FimX could coordinate with another cdG-binding protein to regulate T4P-related processes. Mass spectrometry analysis also identified a number of metabolism-related putative interaction partners. However, interactions between these proteins and FimX require further investigation.

Perhaps it is not surprising that unlike in *P. aeruginosa, A. baylyi* Pil-Chp function occurs independently of cAMP signaling. Previous work showed that the Pil-Chp system in *A. baylyi* regulates T4P positioning rather than T4P activity as it does in *P. aeruginosa* (Ellison *et al*., 2022a). Although cdG binding and cAMP regulation do not appear to be part of the Pil-Chp signaling cascade in *A. baylyi*, it is possible another unidentified signaling molecule may play a role in establishing T4P patterning.

Intriguingly, Kazmierczak *et al* suggested that aberrant FimX may result in mislocalized T4P in *P. aeruginosa* (Kazmierczak *et al*., 2006), though this study was limited by low throughput imaging via electron microscopy. With the development of a broadly applicable labeling methodology for imaging T4P, it is now possible to image T4P phenotypes in diverse species in live cells (including *P. aeruginosa*) without cumbersome sample preparation (Ellison *et al*., 2017, 2018, 2019; Koch *et al*., 2021). Assessing T4P localization in *fimX* mutants in species with polarly localized T4P such as *P. aeruginosa* and *Acinetobacter* pathogens is of significant interest, as it could point to an ancestral function for FimX in regulating T4P machinery placement (Vesel and Blokesch, 2021).

Teasing apart the exact function of FimX and the other Pil-Chp components in Pil-Chp signaling remains challenging as Pil-Chp components influence each other’s localization and stability (Ellison *et al*., 2022a). It is possible that multiple components participate in transient and constant phosphotransfer to maintain interactions that establish linear T4P localization. The localization patterns of individual Pil-Chp components in single deletion mutants may provide some insight into the details of signaling. FimX became cytoplasmically diffuse in nearly 100% of cells in *pilG* or *fimL* mutants, yet a *fimV* deletion resulted in half the cells showing either cytoplasmically diffuse fluorescence or fluorescence localized as a single punctum (Figure 3D). This suggests the nature of FimX interactions with FimV are inherently different from those with PilG or FimL. It is also interesting that deletion of *pilG* or *fimL* causes the other to become cytoplasmically diffuse (Ellison *et al*., 2022a), yet they exhibit different localization patterns in *fimX* mutants (Figure 3E). This may suggest a role for FimX in regulating the interplay between FimL and PilG via ChpA signaling. Analysis of the reciprocal localization of all Pil-Chp component in single deletion mutants is of future interest and will likely provide insight into how these components interact and regulate each other, as will phosphotransfer mutations in ChpA and PilG proteins.

The flagellar chemotaxis system in *Escherichia coli* is known for its ability to rapidly regulate the direction of flagellar rotation in response to changing chemical gradients in the environment (Colin *et al*., 2021). In contrast, the placement of T4P machines is permanent and takes place over much longer time scales. It is unclear why a chemosensory network would regulate generationally long processes, and it is still unclear whether an environmental signal exists. Previous work shows that T4P localization patterns regulate the structure of multicellular communities (Ellison *et al*., 2022a). Because multicellular communities are more resistant to external stresses such as antibiotic exposure, it is probable that a stress signal may trigger changes to T4P patterning to promote cell-cell aggregation. It is also possible that localized signaling between individuals in the community may play a role in regulating T4P patterning to adapt community structure to specific conditions.

Overall, this work highlights an example of how bacteria can adapt accessory proteins to regulate different biological processes. The divergence of FimX away from regulating T4P extension in *A. baylyi* in comparison to its relatives provides a unique platform to dissect the evolution of regulatory networks and their influence on bacterial cell biology.

## MATERIALS AND METHODS

### Bacterial strains, plasmids, and cloning

#### *A. baylyi* strains and culture conditions

*A. baylyi* strain ADP1 was used throughout this study. For a list of strains used throughout, see Supplementary Table 1. *A. baylyi* cultures were grown at 30 °C in lysogeny broth (LB) medium and on agar supplemented with kanamycin (Kan) (50 µg/mL), spectinomycin (60 µg/mL), gentamycin (30 µg/mL), chloramphenicol (30 µg/mL), zeocin (100 µg/mL), and/or apramycin (50 µg/mL) as appropriate.

#### Strain construction in *A. baylyi*

Mutants in *A. baylyi* were made using natural transformation as described previously (Ellison *et al*., 2021, 2022a). Briefly, mutant constructs were made by splicing-by-overlap (SOE) PCR to stich (1) ∼3 kb of the homologous region upstream of the gene of interest, (2) the mutation where appropriate (for example deletion by allelic replacement with an antibiotic [AbR] cassette), and (3) ∼3 kb of the homologous downstream region. For a list of primers used to generate *Acinetobacter* mutants in this study, see Supplementary Table 2. The upstream region was amplified using F1 + R1 primers, and the downstream region was amplified using F2 + R2 primers. All AbR cassettes were amplified with ABD123 (ATTCCGGGGATCCGTCGAC) and ABD124 (TGTAGGCTGGAGCTGCTTC). Tagged proteins were constructed using the primers indicated in Supplementary Table 2. SOE PCR reactions were performed using a mixture of the upstream and downstream regions, and middle region where appropriate using F1 + R2 primers. SOE PCR products were added with 50 µL of overnight-grown culture to 450 µL of LB in 2-mL round-bottom microcentrifuge tubes (USA Scientific) and grown at 30 °C rotating on a roller drum for 3–5 h. For AbR-constructs, transformants were serially diluted and plated on LB and LB + antibiotic.

Complementation strains were constructed by placing the gene of interest under a constitutive *P_tac_* promoter at the *vanAB* locus as described previously (Ellison *et al*., 2021). First, the *vanAB* locus containing the Kan^R^ and the constitutive *P_tac_* promoter published previously (Ellison *et al*., 2021) was amplified using primers CE173 + CE336 to amplify the upstream region. The downstream region was amplified from the same Kan^R^-P*_tac_* construct strain using primers CE406 + CE176. The complementation gene was then amplified using specified primers found in Supplementary Table 2. SOE PCRs and natural transformations were then performed exactly as described above.

Strains containing GFP tagged FimX were constructed at the *vanAB* locus linked to a Kan^R^ cassette for ease of moving the construct into different strains. *P_tac_-fimX-GFP* was constructed using primers CE173 + CE1747 to amplify the upstream region and primers CE2135 + CE176 to amplify the downstream region from a previously constructed strain containing the complementation of FimX at the *vanAB* locus (CE726). Primers CE1555 + CE1556 were used to amplify GFP with designed linker sequences. SOE PCRs and natural transformations were then performed exactly as described above. Strains were confirmed by Sanger sequencing using designed primers indicated in Supplementary Table 2. Strains with FimX tagged at the native locus with either mRuby3 or the 3xFLAG tag were also constructed using SOE PCRs with primers found in Supplementary Table 2.

#### Plasmid construction

Primers used for plasmid construction in this study are listed in Supplementary Table 3. Plasmids used in this study are listed in Supplementary Table 4. Commercially available, chemically competent *Escherichia coli* TOP10 (Invitrogen) was used for plasmid construction and was grown at 37 °C in LB supplemented with 50 μg/mL Kan or 100 μg/mL carbenicillin (Carb), where appropriate, for plasmid maintenance.

For construction of pET28a-derived plasmids, *fimX* from *Pseudomonas aeruginosa* PAO1 or *Acinetobacter baylyi* ADP1 was PCR amplified from their respective genomic DNA and cloned into pET28a plasmid with an N-terminal hexa-histidine tag using Gibson assembly (HiFi DNA Assembly Master Mix; New England Biolabs). The recombinant pET28a::*fimX* vector from each strain was sequence verified using whole plasmid sequencing (Plasmidsaurus).

*A. baylyi* ADP1 *fimX*, *pilB*, *pilZ*, and *chpA* were cloned into BACTH plasmids pUT18C and pUT18 as well as pKT25 and pKNT25, which encodes the T18 and T25 fragments of *Bordetella pertussis* adenylate cyclase at the N- and C-terminal ends, respectively, using Gibson assembly (HiFi DNA Assembly Master Mix; New England Biolabs). All recombinant vectors were sequence verified using whole plasmid sequencing (Plasmidsaurus).

#### Natural transformation assays

Transformation assays in *A. baylyi* were performed exactly as previously described (Ellison *et al*., 2021, 2022a; Ellison and Ellison, 2024). Briefly, strains were grown overnight in LB broth at 30 °C on a roller drum. Then, 50 µL of overnight culture was subcultured into 450 µL of fresh LB medium and at least 50 ng of transforming DNA (tDNA) (a ∼7 kb PCR product containing Δ*pilT*::*spec* amplified using primers CE49 + CE50) was used for all transformation assays except for DNA competition assays which are detailed below. DNA was quantified using a Qubit (ThermoFisher) following standard Qubit protocols. Reactions were incubated with end-over-end rotation on a roller drum at 30 °C for 5 h and then plated for quantitative culture on LB + antibiotic plates (to quantify transformants) and on plain LB plates (to quantify total viable counts). Data are reported as the transformation frequency, which is defined as the (CFU/mL of transformants) / (CFU/mL of total viable counts).

#### Pilin labeling, imaging, and quantification

Pilin labeling in *A. baylyi* was performed as described previously (Ellison *et al*., 2022a; Ellison and Ellison, 2024). Briefly, 100 µL of overnight cultures was added to 900 µL of fresh LB in a 1.5 mL microcentrifuge tube, and cells were grown at 30 °C rotating on a roller drum for 70 min. Cells were then centrifuged at 18,000 x *g* for 1 min and resuspended in 50 µL of LB before labeling with 25 µg/mL of AlexaFluor488 C5-maleimide (AF488-mal) (ThermoFisher) for 15 min at room temperature. Labeled cells were centrifuged, washed three times with 100 µL of PBS and resuspended in 5–40 µL PBS. Cell bodies were imaged using phase-contrast microscopy while labeled pili were imaged using fluorescence microscopy on a Nikon Ti2-E microscope using a Plan Apo 100X oil immersion objective, a GFP/FITC/Cy2 filter set for pili, a Hamamatsu ORCA-Fusion Gen-III cCMOS camera, and Nikon NIS Elements Imaging Software. Cell numbers and the percent of cells making pili were quantified manually using Fiji. All imaging was performed under 1% agarose pads made with PBS solution.

#### FimX imaging and quantification

100 µL of overnight FimX-mRuby3 culture was added to 900 µL of fresh LB in a 1.5 mL microcentrifuge tube, and cells were grown at 30 °C rotating on a roller drum for 70 min. Cells were then centrifuged at 18,000 x *g* for 1 min and resuspended in 5-40 µL of PBS. Cell bodies were imaged using phase-contrast microscopy while labeled FimX were imaged using fluorescence microscopy on a Nikon Ti2-E microscope using a Plan Apo 100X oil immersion objective, a DSRed/TRITC/Cy3 filter set for labeled FimX, a Hamamatsu ORCA-Fusion Gen-III cCMOS camera, and Nikon NIS Elements Imaging Software. Cell numbers and the localization of FimX were quantified manually using Fiji. All imaging was performed under 1% agarose pads made with PBS solution.

#### Coimmunoprecipitation assays

Coimmunoprecipitation experiments were performed as described previously (Ellison *et al*., 2021), with some differences. Cultures of cells were grown shaking overnight either in 3 ml LB in 14 ml culture tubes at 30 °C or in 50 ml LB in 250 ml volume flasks at 30 °C. Overnight cultures of cells grown in tubes in LB medium were diluted by 1/10 into fresh LB for a total volume 50 mL in 250 mL volume flasks. Then 50 mL cultures were grown to exponential growth phase by shaking for 1.5 h at 30 °C. Overnight cultures grown shaking overnight were not placed into exponential. The total culture volume was then harvested at 10,000 × *g* for 10 min at room temperature, and the supernatant was removed. Cell pellets were resuspended in 2 ml of Buffer 1 (50 mM Tris-HCl pH 7.4, 150 mM NaCl, 1 mM EDTA) and transferred to 2 ml volume microcentrifuge tubes and centrifuged at 10,000 × *g* for 3 min. Cells were washed once more with 2 ml of Buffer 1, and washed pellets were resuspended in 1 ml of Buffer 2 (50 mM Tris-HCl pH 7.4, 150 mM NaCl, 1 mM EDTA, 10 mM MgCl_2_, 0.1% Triton X-100, 2% v/v glycerol). To lyse cells, 30 units of DNase I (New England Biolabs) and 10 µl of concentrated protease inhibitor cocktail (Sigma) (one pellet dissolved in 500 µl of ddH2O) were added to cell suspensions and sonicated using a 418 ultrasonic cell disruptor microtip probe (Misonix). Cell debris was removed by centrifugation at 10,000 × *g* for 5 min at 4 °C. 50 µl of cell lysates was added to a fresh 1.7 ml centrifuge tube and set aside for input analysis. 50 µl aliquots of α-FLAG magnetic bead slurry (Sigma) in 1.5 ml microcentrifuge tubes were washed three times with 1 ml of Buffer 2 using a magnetic collection stand. Up to 1 ml of cell lysates was added to washed magnetic beads and subjected to end-over-end rotation at 4 °C for 2 h. Beads were then washed three times with 0.5 ml of Buffer 2, with 10 min incubations in Buffer 2 at 4 °C between each wash step. Beads were briefly washed a 4th time with 0.5 ml Buffer 2. To elute proteins from α-FLAG beads, 100 µl of elution buffer (150 µg/ml 3X-FLAG peptide, Sigma, in Buffer 2) was added and samples were subjected to end-over-end rotation at 4 °C for 30 min. Eluates were mixed with 4X SDS running buffer (40% v/v glycerol, 250 mM Tris-HCl pH 6.8, 0.8% w/v bromophenol blue, 20% v/v β-mercaptoethanol, 8% w/v SDS) and heated to 100 °C for 10–15 min before separation on a 4–20% pre-cast SDS-PAGE gel (BioRad) for ∼15 min so that the total protein was contained in the top ∼1 cm of the gel. Using a new razor blade, the gel was cut to contain only the samples and the protein ladder. Then, a western blot analysis was performed on the gel according to the protocol below.

#### Western blotting of coimmunoprecipitation assay products

After proteins were separated on a 4–20% pre-cast polyacrylamide gel (Biorad) by SDS electrophoresis according to the coimmunoprecipitation protocol previously described, protein bands were electrophoretically transferred to a nitrocellulose membrane, and probed with 1:5000 dilution of mouse monoclonal α-FLAG antibodies (Sigma) and a 1:2500 dilution of mouse monoclonal α-GFP (Roche Diagnostics) primary antibodies. Blots were washed and then probed with a 1:10,000 dilution of goat α-mouse antibody conjugated to horse radish peroxidase (Sigma). Blots were then incubated with SuperSignal West Pico PLUS Chemiluminescence substrate (ThermoFisher) and then imaged using a BioRad Chemidoc imaging system.

#### Mass spectrometry

Mass spectrometry analysis was performed as done previously (Ellison *et al*., 2022a). In-gel digestion of protein bands using trypsin was performed as in Shevchenko *et al*. (Shevchenko *et al*., 2007). Trypsin digested samples were dried completely in a SpeedVac and resuspended with 20 µl of 0.1% formic acid pH 3 in water. 2 µl (∼360 ng) was injected per run using an Easy-nLC 1200 UPLC system. Samples were loaded directly onto a 45 cm long 75 µm inner diameter nano capillary column packed with 1.9 µm C18-AQ resin (Dr. Maisch, Germany) mated to metal emitter in-line with an Orbitrap Fusion Lumos (Thermo Scientific, USA). Column temperature was set at 45 °C and two-hour gradient method with 300 nl per minute flow was used. The mass spectrometer was operated in data dependent mode with the 120,000 resolution MS1 scan (positive mode, profile data type, AGC gain of 4e5, maximum injection time of 54 s and mass range of 375-1500 m/z) in the Orbitrap followed by HCD fragmentation in ion trap with 35% collision energy. Dynamic exclusion list was invoked to exclude previously sequenced peptides for 60 s and maximum cycle time of 3 s was used. Peptides were isolated for fragmentation using quadrupole (1.2 m/z isolation window). Ion-trap was operated in Rapid mode.

Raw files were searched using Byonic (Bern *et al*., 2012) and Sequest HT algorithms (Eng *et al*., 1994) within the Proteome Discoverer 2.2 suite (Thermo Scientific, USA). 10 ppm MS1 and 0.4 Da MS2 mass tolerances were specified. Carbamidomethylation of cysteine was used as fixed modification, oxidation of methionine, deamidation of asparagine and glutamine were specified as dynamic modifications. Pyro glutamate conversion from glutamic acid and glutamine are set as dynamic modifications at peptide N-terminus. Acetylation was specified as dynamic modification at protein N-terminus. Trypsin digestion with maximum of two missed cleavages were allowed. Files were searched against UP000000430 *Acinetobacter baylyi* database downloaded from Uniprot.org.

Scaffold (version Scaffold_4.11.1, Proteome Software Inc., Portland, OR) was used to validate MS/MS based peptide and protein identifications. Peptide identifications were accepted if they could be established at greater than 95.0% probability by the Scaffold Local FDR algorithm. Protein identifications were accepted if they could be established at greater than 99.9% probability and contained at least 2 identified peptides. Protein probabilities were assigned by the Protein Prophet algorithm (Nesvizhskii *et al*., 2003).

#### FimX Expression and Purification

*E. coli* BL21 CodonPlus (DE3) cells were transformed with pET28a::*fimX* from *P. aeruginosa* or *A. baylyi* and grown in 2 L of LB containing 50 μg/mL of Kan at 37 °C to an OD_600_ of 0.6-0.8. Protein expression was induced by the addition of isopropyl-D-1-thiogalactopyranoside (IPTG) to a final concentration of 0.1 mM for 18 hours at 18°C. Cells were harvested by centrifugation at 7000 x *g* for 15 mins. Pellets were resuspended in 40 mL of binding buffer (50 mM HEPES, pH 8, 300 mM NaCl, 10% v/v glycerol, 30 mM imidazole) in addition to a protease inhibitor cocktail tablet (SIGMA*FAST*™, EDTA-free), and small amounts of powdered lysozyme (Bio Basic) and DNase I (Bio Basic). Cell lysis was performed by passage through an Emulsiflex-c3 homogenizer and insoluble cell debris was clarified by centrifugation at 35,000 x *g* for 45 mins. Filtered supernatant was incubated with 2.5 mL of Ni-NTA agarose resin (Qiagen) pre-equilibrated with binding buffer. The column was washed with 75 mL of the same buffer before elution in 25 mL of elution buffer (binding buffer plus 300 mM imidazole). The sample was concentrated using an Amicon Ultra-15 centrifugal device (Millipore) with 100 kDa molecular weight cut-off at 4000 x *g* in ten-minute intervals and subsequently injected into a HiLoad 16/60 Superdex 200 prep grade gel filtration column (GE Healthcare) pre-equilibrated in SEC buffer (50 mM HEPES, pH 8, 200 mM NaCl, 10% v/v glycerol) for further purification. Fractions containing FimX were assessed by 12% SDS-PAGE gel, concentrated, and flash frozen in liquid nitrogen for storage at -80°C until use.

#### Isothermal Titration Calorimetry

Cyclic di-guanosine monophosphate (cdG) (InvivoGen) (250 μM in the syringe) and *Pa*FimX or *Ab*FimX proteins (25 μM in the cell) were prepared in SEC buffer. Guanosine triphosphate (GTP) (Sigma) (1 mM in the syringe) and FimX proteins (100 μM in the cell) were prepared in SEC buffer plus 10 mM MgCl_2_. Calorimetric titrations were conducted using a MicroCal PEAQ-ITC Automated microcalorimeter (Malvern Panalytical) at 25°C, comprising 19 2-μL injections of 4 s per injection with 120 s intervals between injections. The first titration was 0.4 μL for 0.8 s. The MicroCal PEAQ-ITC software v.1.41 (Malvern Panalytical) was used to subtract the heats of dilution for titrating cdG or GTP into buffer from the sample and to perform curve-fitting using a single-site binding model to derive the dissociation constant.

#### Bacterial Adenylate Cyclase Two-Hybrid (BACTH) Assays

Combinations of the *A. baylyi* FimX and PilB or PilZ or ChpA T18 and T25 fusion proteins were transformed into *E. coli* BTH101 (Euromedex). Overnight cultures of each combination were inoculated with 5 mL of LB supplemented with 50 μg/mL of Kan, 100 μg/mL of Carb, and 0.5 mM of IPTG and grown overnight at 30 °C for 18-20 hours. For the plates showing all combinations or the plates that complemented the quantitative Miller assay, 5 or 10 μL, respectively, of overnight cultures was spotted onto LB agar plate containing 50 μg/mL of Kan, 100 μg/mL of Carb, 0.5 mM of IPTG, and 50 μg/mL of 5-bromo-4-chloro-3-indolyl-β-D-galactopyranoside (X-Gal). Plates were incubated at 30 °C for 48 hours and subsequently at 4 °C for 24 hours prior to imaging. pUT18C::zip and pKT25::zip vector pairs and empty pUT18C and pKT25 vector pairs were used as positive and negative controls, respectively.

A ꞵ-galactosidase Miller assay was performed to quantify the strength of the BACTH interactions. Overnight cultures grown in 5 mL LB supplemented with 50 μg/mL of Kan, 100 μg/mL of Carb, and 0.5 mM of IPTG at 30 °C for 20 hours were normalized to an OD_600_ of 1 in a 1 mL volume and centrifuged at 12,000 rpm for 5 mins. Cell pellets were resuspended in 500 μL of 60 mM Na_2_HPO_4_, 40 mM NaH_2_PO_4_, pH 7, 20 mM KCl, 2 mM MgSO_4_, 0.4 mg/mL sodium deoxycholate, 0.8 mg/mL cetyltrimethylammonium bromide, and 75 mM β-mercaptoethanol and incubated at 30 °C for 30 mins. 200 μL of cellular material was mixed with 600 μL of 60 mM Na_2_HPO_4_, 40 mM NaH_2_PO_4_, pH 7, and 1 mg/mL ortho-nitrophenyl-β-galactoside and incubated at 30 °C for 20 mins. 700 μL of 1 M Na_2_CO_3_ solution was added to stop the reaction and centrifuged for 10 mins. 200 μL of each reaction was measured in a 96-well plate in triplicates at absorbances of 420 nm and 550 nm. ꞵ-galactosidase activity in Miller units was calculated using the following equation: [A_420_ – (1.75 x A_550_)] / (reaction time in minutes x OD_600_ x volume of reaction assayed in mL) x 1000.

#### Western Blotting of BACTH constructs

Overnight cultures grown in 5 mL LB supplemented with 50 μg/mL of Kan, 100 μg/mL of Carb, and 0.5 mM of IPTG at 30 °C for 20 hours were normalized to an OD_600_ of 1 in a 1 mL volume. Cells were pelleted by centrifugation at 15,000 rpm for 5 mins. After removing the supernatant, cell pellets were resuspended in 100 μL of 2× Laemmli buffer, boiled for 20 mins, and separated by SDS-PAGE using 12% gels. Protein bands were transferred to 0.2-µm PVDF membrane (25 V, 1.5 hours) with Towbin buffer (25 mM Tris-HCl, pH 8.6, 192 mM glycine, 20% v/v methanol) and blocked in 5% (w/v) skim milk powder resuspended in Tris buffered saline with Tween-20 (TBS-T; 20 mM Tris-HCl pH 7.6, 150 mM NaCl, 0.05% v/v Tween-20) for 1 hour. Membranes were probed with α-adenylate cyclase toxin mouse monoclonal antibodies (Santa Cruz Biotechnology) at 1:5000 dilution in 1% (w/v) skim milk powder resuspended in TBS-T for 20 hours. Membranes were then washed four times with TBS-T and probed with horseradish peroxidase (HRP)-conjugated goat anti-mouse antibody (Biorad) at 1:3000 dilution in 1% (w/v) skim milk powder resuspended in TBS-T for 1 hour. Membranes were washed three times with TBS-T and developed using SuperSignal West Pico Plus chemiluminescent substrate (ThermoFisher Scientific).

## ACKNOWLEDGMENTS

We thank Ivan Ristic for assistance with cloning the BACTH plasmids. The work was supported in part by Canadian Institutes of Health Research Project grant PJT-169053 to P.L.H. I.Y.Y. was supported by a Natural Sciences and Engineering Research Council PGS-D scholarship and a Hospital for Sick Children Research Institute Restracomp scholarship. P.L.H. was the recipient of a Tier I Canada Research Chair in Structural Biology from 2006-2020. ITC was performed at the Structural & Biophysical Core (SBC) facility at the Hospital for Sick Children with advice from Mario Vargus and Magnus Jorgensen. Courtney Ellison, PhD, is a Damon Runyon-Marilyn and Scott Urdang Breakthrough Scientist supported by the Damon Runyon Cancer Research Foundation (DFS6023). This work was supported by a Hypothesis Fund award and by the National Institutes of Health grant R35GM150916 awarded to C.K.E.

