## Supplemental information for "FimX regulates type IV pilus localization via the Pil-Chp chemosensory system in *Acinetobacter baylyi*"

**Supplementary Table 1. Bacterial strains used throughout this study.**

| Strain CE # | Strain name in manuscript | Genotype | Figure ('S' denotes Supplementary figure) | Source |
|---|---|---|---|---|
| CE100 | Parent | ADP1 *comP*$^{T129C}$ | 1C, 2A, 2B, 2C, 4B, 5A, 5B, 5C, 6A | Ellison et al. 2021 |
| CE1973 | FimX-GFP | ADP1 *comP*$^{T129C}$ *ΔvanAB::kan*$^{R}$, *P*$_{tac}$*-fimX-GFP ΔfimX::chlor*$^{R}$ | 1C, 6A | This study |
| CE2021 | FimX-GFP FLAG-PilB | ADP1 *comP*$^{T129C}$ *3xFLAG-pilB ΔvanAB::kan*$^{R}$, *P*$_{tac}$*-fimX-GFP ΔfimX::chlor*$^{R}$ | 1C | This study |
| CE2023 | FimX-GFP FLAG-PilZ | ADP1 *comP*$^{T129C}$ *3xFLAG-pilZ ΔvanAB::kan*$^{R}$, *P*$_{tac}$*-fimX-GFP ΔfimX::chlor*$^{R}$ | 1C | This study |
| CE358 | *ΔcomP* | ADP1 *ΔcomP::kan*$^{R}$ | 2A | Ellison et al. 2021 |
| CE712 | *ΔfimX* | ADP1 *comP*$^{T129C}$ *ΔfimX::chlor*$^{R}$ | 2A, 2C | This study |
| CE726 | *ΔfimX P*$_{tac}$*-fimX* | ADP1 *comP*$^{T129C}$ *ΔfimX::chlor*$^{R}$ *ΔvanAB::kan*$^{R}$, *P*$_{tac}$*-fimX* | 2A, 2C | This study |
| CE317 | Parent | ADP1 *comP*$^{T129C}$ *ΔpilT::spec*$^{R}$ | 2B, 2C, 2D, S4 | Ellison et al. 2021 |
| CE728 | *ΔfimX* | ADP1 *comP*$^{T129C}$ *ΔfimX::chlor*$^{R}$ *ΔpilT::spec*$^{R}$ | 2B, 2C, 2D, S4 | This study |
| CE1775 | *ΔfimX P*$_{tac}$*-fimX* | ADP1 *comP*$^{T129C}$ *ΔfimX::chlor*$^{R}$ *ΔvanAB::kan*$^{R}$, *P*$_{tac}$*-fimX ΔpilT::spec*$^{R}$ | 2B, 2C, 2D | This study |
| CE1981 | *ΔfimX P*$_{tac}$*-fimX-GFP* | ADP1 *comP*$^{T129C}$ *ΔvanAB::kan*$^{R}$, *P*$_{tac}$*-fimX-GFP ΔfimX::chlor*$^{R}$ *ΔpilT::spec*$^{R}$ | S4 | This study |
| CE2131 | *fimX-mRuby3* | ADP1 *comP*$^{T129C}$ *fimX-mRuby3 ΔpilT::spec*$^{R}$ | S4 | This study |
| CE2134 | *fimX-3xFLAG* | ADP1 *comP*$^{T129C}$ *fimX-3xFLAG ΔpilT::spec*$^{R}$ | S4 | This study |
| CE765 | FimX-mRuby3, Parent | ADP1 *comP*$^{T129C}$ *fimX-mRuby3* | 3B, 3C, 3D | This study |
| CE789 | *ΔpilG* | ADP1 *comP*$^{T129C}$ *fimX-mRuby3 ΔpilG* | 3D | This study |
| CE787 | *ΔfimL* | ADP1 *comP*$^{T129C}$ *fimX-mRuby3 ΔfimL::chlor*$^{R}$ | 3D | This study |
| CE786 | *ΔfimV* | CE786: ADP1 *comP*$^{T129C}$ *fimX-mRuby3 ΔfimV::kan*$^{R}$ | 3D | This study |
| CE17 | PilG + | ADP1 *comP*$^{T129C}$ *pilG-mRuby3* | 3E | Ellison et al. 2022a |
| CE468 | PilG - | ADP1 *comP*$^{T129C}$ *pilG-mRuby3 ΔfimX::kan*$^{R}$ | 3E | This study |
| CE251 | FimL + | ADP1 *comP*$^{T129C}$ *fimL-mCherry* | 3E | Ellison et al. 2022a |
| CE504 | FimL - | ADP1 *comP*$^{T129C}$ *fimL-mCherry ΔfimX::kan*$^{R}$ | 3E | This study |
| CE745 | FimV + | ADP1 *comP*$^{T129C}$ *fimV-mRuby3* | 3E | Ellison et al. 2022a |
| CE759 | FimV - | ADP1 *comP*$^{T129C}$ *fimV-mRuby3 ΔfimX::kan*$^{R}$ | 3E | This study |
|  | PaFimX | *Pseudomonas aeruginosa* PAO1 | 4B | M. R. Parsek |
| CE1883 | *ΔcyaB* | ADP1 *comP*$^{T129C}$ *ΔcyaB::apr*$^{R}$ | 5A, 5B | This study |

| | | | | |
|---|---|---|---|---|
| CE34 | ∆*vfr* | ADP1 *comP*$^{T129C}$ ∆*crp::kan*$^{R}$ | 5A, 5B | This study |
| CE2020 | FimX-GFP ChpA-FLAG | ADP1 *comP*$^{T129C}$ *chpA-3xFLAG* ∆*vanAB::kan*$^{R}$, *P*$_{tac}$-*fimX-GFP* ∆*fimX::chlor*$^{R}$ | 6A | This study |
| *E. coli* BTH101 (BACTH strain) | | F−*cya-99 araD139 galE15 galK16 rpsL1 (Str*$^{R}$*) hsdR2 mcrA1 mcrB1NEB5α pNPTS138::ΔcpaF* | S1, S2, 1A, 1B, 6B, 6C, S5 | Euromedex |
| *E. coli* BL21 CodonPlus (DE3) | | F-, *ompT hsdS(rB-mB-) dcm+Tet*$^{R}$*galλ (DE3) endA [argU proLCam*$^{R}$*]* | Protein expression strain | Stratagene |
| *E. coli* TOP10 | | F-*mcrA∆(mrr-hsdRMS-mcrBC) Φ80lacZ∆M15 ∆lacX74∆araD139∆(ara leu)7697 galU galK rpsL endA 1nupG, Str*$^{R}$ | Cloning Strain | Invitrogen |

**Supplementary Table 2. Primers used for *Acinetobacter* strain construction in this study.**

| **Primer #** | **Sequence 5' → 3' (overlapping regions in lower case, 3xFLAG tag in bold)** | **Description** |
|---|---|---|
| 1149 | CGAAGTTTTACACGGTTGAATGCTTGAC | fimX F1 |
| 1150 | gtcgacggatccccggaatACGCTTTATTTTTTTTGCTAGCAACCCG | AbR replace fimX R1 |
| 1151 | gaagcagctccagcctacaGGTAAATTAGACCGTCTTGTTGATGTGCA | AbR relace fimX F2 |
| 1152 | TTGGTCTTGTAAAGTCGGGTAAGCC | fimX R2 (and fimX Cterm R2) |
| 1147 | GCCATGAACAACAGCAATCTATTAGTATTTATGTC | fimX confirmation F |
| 1148 | GCCACCCATTCTATACTGGTAAACAAAAGAA | fimX confirmation R |
| 233 | ATTCCGGGGATCCGTCGAC | AbR cassette F |
| 234 | TGTAGGCTGGAGCTGCTTC | AbR cassette R |
| 317 | GCAAACCACAAACATAATGTTTGAAATCC | vanAB::kan F1 for ectopic expression products |
| 176 | CCAAGACTATAAATAATCGACATGATCAATTTTAA | vanAB R2 for ectopic expression constructs |
| 49 | GTACTCATCTCGCATATTCAGGAAATG | pilT F1 (and tDNA F1) |
| 50 | CGCAAGTTCAACAGTCCTACCAA | pilT R2 (and tDNA R2) |
| 19 | CTTTAAAGTGATTCATTGACAGGGTTC | pilT confirmation F |
| 20 | GTAATTTCACCGTTAATTTTTAAAGATGGTTCT | pilT confirmation F |
| 1214 | CTTTATCAGTGCAGTTTTGAATGGGC | fimX Cterm F1 |
| 1747 | tccaccacttccacctgcTTGATCCTGCACATCAACAAGACG | fimX Cterm linker R1 |
| 1748 | gcaggtggagcaggtggaTAACCATTGAACTCCAGACAAAAAAAACG | fimX Cterm linker F2 |
| 1777 | gcaggtggaagtggtg**gagattataaagatcatgatggtgattataaagatcatgatattgattataaa**gatgatgatgataaagcaggtggagcaggtgga | 3xFLAG-linker F |
| 1778 | tccacctgctccacctgc**tttatcatcatcatctttataatcaatatcatgatctttataatcaccatcatgatctttataatc**tccaccacttccacctgc | 3xFLAG-linker R |
| 1555 | gcaggtggaagtggtgga | Fluorescent tag linker F |
| 1556 | tccacctgctccacctgc | Fluorescent tag linker R |
| 31 | GCTGTTTGATTTTATCCAGAACCTTG | pilG F1 |
| 34 | GCGCTGTAATTACAATAGTGTTCGGA | pilG R2 |

| | | |
|---|---|---|
| 29 | AATATCTTGTCGCCGATTCATTGAAA | pilG confirmation F |
| 30 | GCAAGTGTTACCCCATCTGCG | pilG confirmation R |
| 735 | GTGCACTGGGGATATCTGCACT | fimL F1 |
| 738 | TGCTAACAGAATTGTATGTGTCTACGCC | fimL R2 |
| 733 | GCTATCAGGAAAAAAATGGCGAGC | fimL confirmation F |
| 734 | TTCAGGAATTGTTTCATCAGCGCC | fimL confirmation R |
| 1525 | GCCCCGTAATAGTAATTCAAAGTACATGG | fimV F1 |
| 1528 | GTATAAATGAAGAACATACAAGGAAGGGCG | fimV R2 |
| 1523 | GTTGCTCAGAAAGGGTATGACAAG | fimV confirmation F |
| 1524 | GGGGCTTAAATGTCTGAAAAGGC | fimV confirmation R |
| 189 | TCCTGTTCTTTTGGTCGCAGAA | cyaB F1 |
| 3166 | gtcgacggatccccggaatCCTATCCTCCGCAACACAGTT | AbR replace cyaB R1 |
| 3167 | gaagcagctccagcctacaAAAAGCCTTCTATAAATAGAAGGCCTG | AbR replace cyaB F2 |
| 192 | CTCGATGGTTTAGAGATGGATTATGTCA | cyaB F2 |
| 187 | CCATATTCCAATGATGTATTGGCACA | cyaB confirmation F |
| 188 | TGAGCCTGATTCACGTTTATCTTGT | cyaB confirmation R |
| 183 | CGCAATAGTTTTGGCTCTTCATTTT | Crp (Vfr PAO1) F1 |
| 235 | gtcgacggatccccggaatTACTTAATGTTTCTGAAGAGGATAATAAGATCG C | AbR replace crp (Vfr PAO1) R1 |
| 236 | gaagcagctccagcctacaCTTAATCGGTATGTCGTAATCTTAAGAGAGC | AbR replace crp (Vfr PAO1) F2 |
| 186 | GCCGTCATTTGGGTACAGTCG | Crp (Vfr PAO1) R2 |
| 173 | TGACGGTTATCTACCTCTCCAATTTTC | vanAB F1 for ectopic expression products |
| 2135 | gcaggtggagcaggtggaTAAGAAGCAGCTCCAGCCTACA | vanAB locus Cterm linker F2 |
| 457 | ggttctggcaaatattctgaaatgagct | vanAB Ptac-confirmation F |
| 172 | TAGTGAATCGTCAAGCTGGACG | vanAB confirmation R |
| 1231 | GCCAATGCCTGGAATGTAAGTACTC | fimX Cterm confirmation F |

**Supplementary Table 3. Primers used in plasmid construction in this study.**

| **Primer** | **Sequence 5' → 3' (Sequences of homologous regions for Gibson assembly are bolded)** |
|---|---|
| pET28fimX-PAO1-FI | **GCGCGGCAGCCAT**ATGGCCATCGAAAAGAAAACCATC |
| pET28fimX-PAO1-RI | **CTTGTCGACGGAGCTCGAATT**CTCATTCGTCTCCCGAGGAGAAG |
| pET28fimX-PAO1-FV | **CTTCTCCTCGGGAGACGAATGA**GAATTCGAGCTCCGTCGACAAG |
| pET28fimX-PAO1-RV | **GATGGTTTTCTTTTCGATGGCCAT**ATGGCTGCCGCGC |
| pET28fimX-ADP1-FI | **GCGCGGCAGCCAT**ATGAGAAACGGGTTGCTAGCA |
| pET28fimX-ADP1-RI | **CTTGTCGACGGAGCTCGAATTC**TTATTGATCCTGCACATCAACAAGACG |
| pET28fimX-ADP1-FV | **CGTCTTGTTGATGTGCAGGATCAATAA**GAATTCGAGCTCCGTCGACAAG |
| pET28fimX-ADP1-RV | **TGCTAGCAACCCGTTTCTCAT**ATGGCTGCCGCGC |
| pUT18fimX-FI | **CAAGCTTGCATGCCTGCAGGTCGACT**GTGAGAAACGGGTTGCTAGCA |
| pUT18fimX-RI | **GCTGGCGGCTGAATTCGAGCTCGG**TTGATCCTGCACATCAACAAGACG |
| pUT18fimX-FV | **CGTCTTGTTGATGTGCAGGATCAA**CCGAGCTCGAATTCAGCCGCCAGC |
| pUT18fimX-RV | **TGCTAGCAACCCGTTTCTCAC**AGTCGACCTGCAGGCATGCAAGCTTG |
| pUT18CfimX-FI | **TGGAACGCCACTGCAGGTCGACT**GTGAGAAACGGGTTGCTAGCA |
| pUT18CfimX-RI | **ATGAATTCGAGCTCGGTACCCGGGG**TTATTGATCCTGCACATCAACAAGACG |
| pUT18CfimX-FV | **CGTCTTGTTGATGTGCAGGATCAATAA**CCCCGGGTACCGAGCTCGAATTCAT |

| | |
|---|---|
| pUT18CfimX-RV | **TGCTAGCAACCCGTTTCTCAC**AGTCGACCTGCAGTGGCGTTCCA |
| pKNT25fimX-FI | **CAAGCTTGCATGCCTGCAGGTCGACT**GTGAGAAACGGGTTGCTAGCA |
| pKNT25fimX-RI | **GCTGCATGGTCATTGAATTCGAGCTCGG**TTGATCCTGCACATCAACAAGACG |
| pKNT25fimX-FV | **CGTCTTGTTGATGTGCAGGATCAA**CCGAGCTCGAATTCAATGACCATGCAGC |
| pKNT25fimX-RV | **TGCTAGCAACCCGTTTCTCAC**AGTCGACCTGCAGGCATGCAAGCTTG |
| pKT25fimX-FI | **GGGCTGCAGGGTCGACTCTAGAG**GTGAGAAACGGGTTGCTAGCA |
| pKT25fimX-RI | **CGACGTTGTAAAACGACGGCCGAATTCT**TTATTGATCCTGCACATCAACAAGACG |
| pKT25fimX-FV | **CGTCTTGTTGATGTGCAGGATCAATAA**AGAATTCGGCCGTCGTTTTACAACGTCG |
| pKT25fimX-RV | **TGCTAGCAACCCGTTTCTCAC**CTCTAGAGTCGACCCTGCAGCCC |
| pUT18pilB-FI | **CAAGCTTGCATGCCTGCAGGTCGACT**ATGTCAGCATTTACAACAC |
| pUT18pilB-RI | **GCTGGCGGCTGAATTCGAGCTCGG**TTCACTGGTTACACGATTAATTTCCTGTAAC |
| pUT18pilB-FV | **GTTACAGGAAATTAATCGTGTAACCAGTGAA**CCGAGCTCGAATTCAGCCGCCAGC |
| pUT18pilB-RV | **GTGTTGTAAATGCTGACAT**AGTCGACCTGCAGGCATGCAAGCTTG |
| pUT18CpilB-FI | **TGGAACGCCACTGCAGGTCGACT**ATGTCAGCATTTACAACAC |
| pUT18CpilB-RI | **ATGAATTCGAGCTCGGTACCCGGGG**TTATTCACTGGTTACACGATTAATTTCCTG |
| pUT18CpilB-FV | **CAGGAAATTAATCGTGTAACCAGTGAATAA**CCCCGGGTACCGAGCTCGAATTCAT |
| pUT18CpilB-RV | **GTGTTGTAAATGCTGACAT**AGTCGACCTGCAGTGGCGTTCCA |
| pKNT25pilB-FI | **CAAGCTTGCATGCCTGCAGGTCGACT**ATGTCAGCATTTACAACAC |
| pKNT25pilB-RI | **GCTGCATGGTCATTGAATTCGAGCTCGG**TTCACTGGTTACACGATTAATTTCCTGTAAC |
| pKNT25pilB-FV | **GTTACAGGAAATTAATCGTGTAACCAGTGAA**CCGAGCTCGAATTCAATGACCATGCAGC |
| pKNT25pilB-RV | **GTGTTGTAAATGCTGACAT**AGTCGACCTGCAGGCATGCAAGCTTG |
| pKT25pilB-FI | **GGGCTGCAGGGTCGACTCTAGAG**ATGTCAGCATTTACAACAC |
| pKT25pilB-RI | **CGACGTTGTAAAACGACGGCCGAATTCT**TTATTCACTGGTTACACGATTAATTTCCTG |
| pKT25pilB-FV | **CAGGAAATTAATCGTGTAACCAGTGAATAA**AGAATTCGGCCGTCGTTTTACAACGTCG |
| pKT25pilB-RV | **GTGTTGTAAATGCTGACAT**CTCTAGAGTCGACCCTGCAGCCC |
| pUT18pilZ-FI | **CAAGCTTGCATGCCTGCAGGTCGACT**ATGGATTCAAGAATCAATGGTGGTCTG |
| pUT18pilZ-RI | **GCTGGCGGCTGAATTCGAGCTCGG**CATGGTAAAGTTTGAACGGTCAGAGC |
| pUT18pilZ-FV | **GCTCTGACCGTTCAAACTTTACCATG**CCGAGCTCGAATTCAGCCGCCAGC |
| pUT18pilZ-RV | **CAGACCACCATTGATTCTTGAATCCAT**AGTCGACCTGCAGGCATGCAAGCTTG |
| pUT18CpilZ-FI | **TGGAACGCCACTGCAGGTCGACT**ATGGATTCAAGAATCAATGGTGGTCTG |
| pUT18CpilZ-RI | **ATGAATTCGAGCTCGGTACCCGGGG**TTACATGGTAAAGTTTGAACGGTCAGAGC |
| pUT18CpilZ-FV | **GCTCTGACCGTTCAAACTTTACCATGTAA**CCCCGGGTACCGAGCTCGAATTCAT |
| pUT18CpilZ-RV | **CAGACCACCATTGATTCTTGAATCCAT**AGTCGACCTGCAGTGGCGTTCCA |
| pKNT25pilZ-FI | **CAAGCTTGCATGCCTGCAGGTCGACT**ATGGATTCAAGAATCAATGGTGGTCTG |
| pKNT25pilZ-RI | **GCTGCATGGTCATTGAATTCGAGCTCGG**CATGGTAAAGTTTGAACGGTCAGAGC |
| pKNT25pilZ-FV | **GCTCTGACCGTTCAAACTTTACCATG**CCGAGCTCGAATTCAATGACCATGCAGC |
| pKNT25pilZ-RV | **CAGACCACCATTGATTCTTGAATCCAT**AGTCGACCTGCAGGCATGCAAGCTTG |
| pKT25pilZ-FI | **GGGCTGCAGGGTCGACTCTAGAG**ATGGATTCAAGAATCAATGGTGGTCTG |
| pKT25pilZ-RI | **CGACGTTGTAAAACGACGGCCGAATTCT**TTACATGGTAAAGTTTGAACGGTCAGAGC |
| pKT25pilZ-FV | **GCTCTGACCGTTCAAACTTTACCATGTAA**AGAATTCGGCCGTCGTTTTACAACGTCG |
| pKT25pilZ-RV | **CAGACCACCATTGATTCTTGAATCCAT**CTCTAGAGTCGACCCTGCAGCCC |
| pUT18chpA-FI | **CAAGCTTGCATGCCTGCAGGTCGACT**ATGGATGCACAAGTAAATCATCTTATAG |
| pUT18chpA-RI | **GCTGGCGGCTGAATTCGAGCTCGG**TTGGTTTTGGCTATGTTTAATCGC |
| pUT18chpA-FV | **GCGATTAAACATAGCCAAAACCAA**CCGAGCTCGAATTCAGCCGCCAGC |
| pUT18chpA-RV | **CTATAAGATGATTTACTTGTGCATCCAT**AGTCGACCTGCAGGCATGCAAGCTTG |
| pUT18CchpA-FI | **TGGAACGCCACTGCAGGTCGACT**ATGGATGCACAAGTAAATCATCTTATAG |
| pUT18CchpA-RI | **ATGAATTCGAGCTCGGTACCCGGGG**TTATTGGTTTTGGCTATGTTTAATCGC |
| pUT18CchpA-FV | **GCGATTAAACATAGCCAAAACCAATAA**CCCCGGGTACCGAGCTCGAATTCAT |
| pUT18CchpA-RV | **CTATAAGATGATTTACTTGTGCATCCAT**AGTCGACCTGCAGTGGCGTTCCA |
| pKNT25chpA-FI | **CAAGCTTGCATGCCTGCAGGTCGACT**ATGGATGCACAAGTAAATCATCTTATAG |

| | |
|---|---|
| pKNT25chpA-RI | **GCTGCATGGTCATTGAATTCGAGCTCGG**TTGGTTTTGGCTATGTTTAATCGC |
| pKNT25chpA-FV | **GCGATTAAACATAGCCAAAACCAA**CCGAGCTCGAATTCAATGACCATGCAGC |
| pKNT25chpA-RV | **CTATAAGATGATTTACTTGTGCATCCAT**AGTCGACCTGCAGGCATGCAAGCTTG |
| pKT25chpA-FI | **GGGCTGCAGGGTCGACTCTAGAG**ATGGATGCACAAGTAAATCATCTTATAG |
| pKT25chpA-RI | **CGACGTTGTAAAACGACGGCCGAATTCT**TTATTGGTTTTGGCTATGTTTAATC |
| pKT25chpA-FV | **GCGATTAAACATAGCCAAAACCAATAA**AGAATTCGGCCGTCGTTTTACAACGTCG |
| pKT25chpA-RV | **CTATAAGATGATTTACTTGTGCATCCAT**CTCTAGAGTCGACCCTGCAGCCC |

**Supplementary Table 4. Plasmids used in this study.**

| Plasmid name | Description | Source |
|---|---|---|
| pET28a | IPTG-inducible expression vector encoding N-terminal hexa-histidine tag, a thrombin cleavage site, and an optional C-terminal hexahistidine tag, Kan$^R$ | Novagen |
| pET28a::*fimX* | pET28a with WT *P. aeruginosa* PAO1 or *A. baylyi* ADP1 *fimX* fused to an N-terminal hexa-histidine tag; Kan$^R$ | This study |
| pUT18 | Encodes T18 fragment of *B. pertussis* adenylate cyclase toxin on C-terminus, Amp$^R$ | Euromedex |
| pUT18C | Encodes T18 fragment of *B. pertussis* adenylate cyclase toxin on N-terminus, Amp$^R$ | Euromedex |
| pKNT25 | Encodes T25 fragment of *B. pertussis* adenylate cyclase toxin on C-terminus, Kan$^R$ | Euromedex |
| pKT25 | Encodes T25 fragment of *B. pertussis* adenylate cyclase toxin on N-terminus, Kan$^R$ | Euromedex |
| pUT18C::zip | BACTH positive control, pUT18C containing sequence coding for the leucine zipper region of yeast GCN4 protein, Amp$^R$ | Euromedex |
| pKT25::zip | BACTH positive control, pKT25 containing sequence coding for the leucine zipper region of yeast GCN4 protein, Kan$^R$ | Euromedex |
| pUT18::*fimX* | *A. baylyi* ADP1 *fimX* cloned into the SalI and SacI sites of pUT18, Amp$^R$ | This study |
| pUT18C::*fimX* | *A. baylyi* ADP1 *fimX* cloned into the SalI and SmaI sites of pUT18C, Amp$^R$ | This study |
| pKNT25::*fimX* | *A. baylyi* ADP1 *fimX* cloned into the SalI and SacI sites of pKNT25, Kan$^R$ | This study |
| pKT25::*fimX* | *A. baylyi* ADP1 *fimX* cloned into the XbaI and EcoRI sites of pKT25, Kan$^R$ | This study |
| pUT18::*pilB* | *A. baylyi* ADP1 *pilB* cloned into the SalI and SacI sites of pUT18, Amp$^R$ | This study |
| pUT18C::*pilB* | *A. baylyi* ADP1 *pilB* cloned into the SalI and SmaI sites of pUT18C, Amp$^R$ | This study |
| pKNT25::*pilB* | *A. baylyi* ADP1 *pilB* cloned into the SalI and SacI sites of pKNT25, Kan$^R$ | This study |
| pKT25::*pilB* | *A. baylyi* ADP1 *pilB* cloned into the XbaI and EcoRI sites of pKT25, Kan$^R$ | This study |
| pUT18::*pilZ* | *A. baylyi* ADP1 *pilZ* cloned into the SalI and SacI sites of pUT18, Amp$^R$ | This study |
| pUT18C::*pilZ* | *A. baylyi* ADP1 *pilZ* cloned into the SalI and SmaI sites of pUT18C, Amp$^R$ | This study |
| pKNT25::*pilZ* | *A. baylyi* ADP1 *pilZ* cloned into the SalI and SacI sites of pKNT25, Kan$^R$ | This study |

| | | |
|---|---|---|
| pKT25::*pilZ* | *A. baylyi* ADP1 *pilZ* cloned into the XbaI and EcoRI sites of pKT25, $Kan^R$ | This study |
| pUT18::*chpA* | *A. baylyi* ADP1 *chpA* cloned into the SalI and SacI sites of pUT18, $Amp^R$ | This study |
| pUT18C::*chpA* | *A. baylyi* ADP1 *chpA* cloned into the SalI and SmaI sites of pUT18C, $Amp^R$ | This study |
| pKNT25::*chpA* | *A. baylyi* ADP1 *chpA* cloned into the SalI and SacI sites of pKNT25, $Kan^R$ | This study |
| pKT25::*chpA* | *A. baylyi* ADP1 *chpA* cloned into the XbaI and EcoRI sites of pKT25, $Kan^R$ | This study |

### Supplemental Figure S1

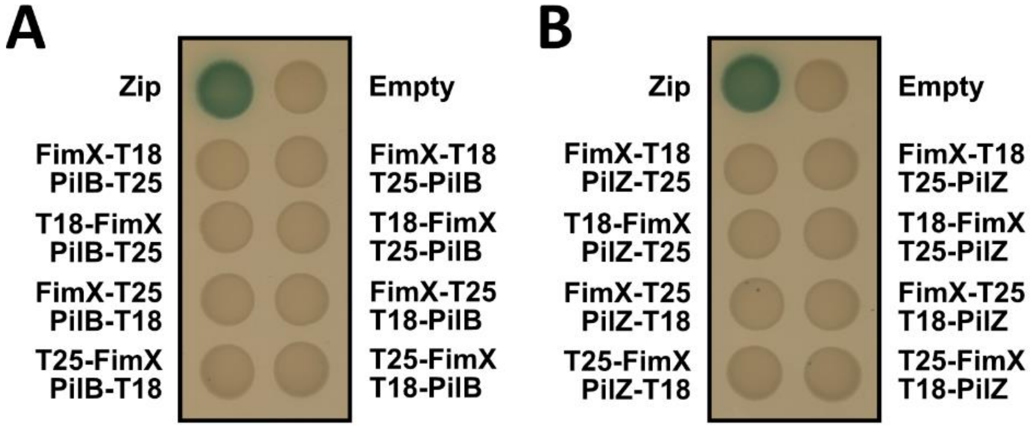


Supplementary Figure S1. FimX does not directly interact with PilB nor PilZ. A-B. All combinations of BACTH interactions between FimX and PilB/PilZ fused to the N- or C-terminus of the T25 and T18 domains of adenylate cyclase following incubation at 30 °C for 48 hours and subsequently at 4 °C for 24 hours on LB agar plates containing X-Gal. Leucine zipper motif (zip) fused to T18 and T25 was used as positive control while empty vectors of T18 and T25 alone (empty) was the negative control.

### Supplemental Figure S2

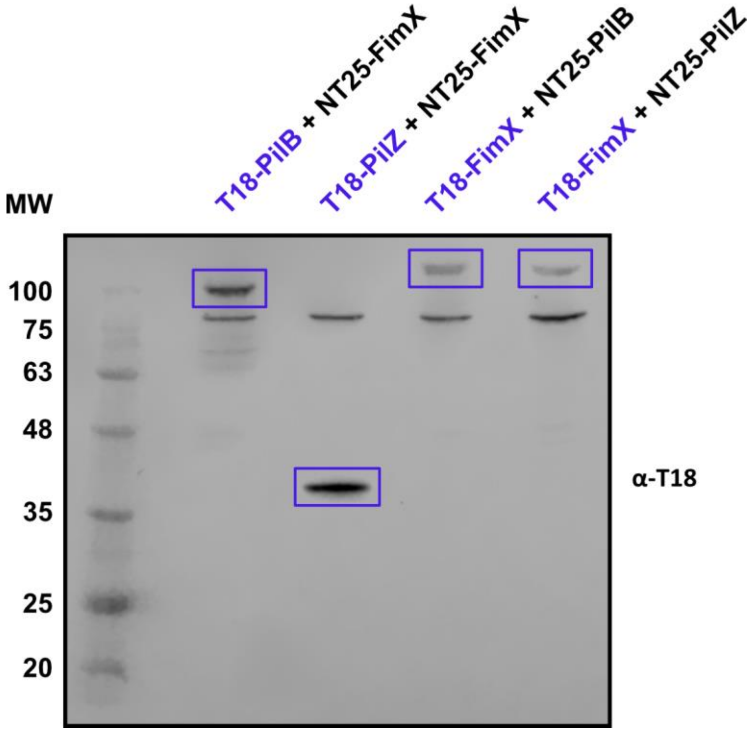


Supplementary Figure S2. PilB, PilZ, and FimX expression in the BACTH system. Western blotting of whole cell lysates from *E. coli* BTH101 cells co-expressing combinations of PilB/PilZ/FimX fused N-terminally to the T18 and T25 domains of adenylate cyclase and probed using a monoclonal antibody against the T18 fragment of adenylate cyclase. Bands corresponding to each fusion protein are boxed in purple. The molecular weight (MW) ladder has units in kilodaltons.

### Supplemental Figure S3

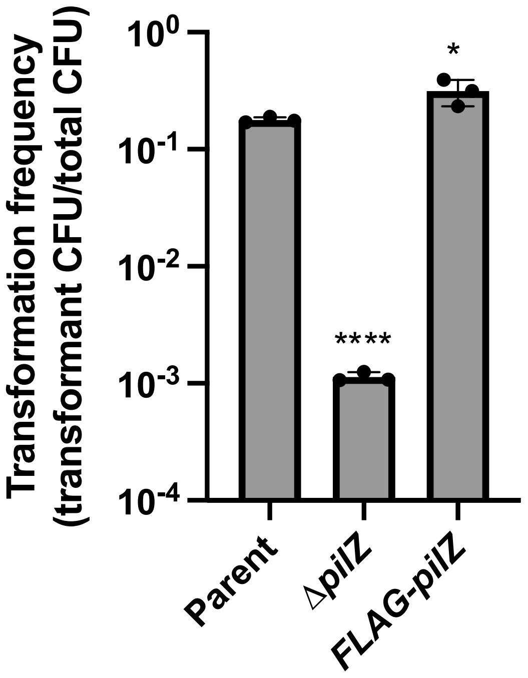


Supplemental Figure S3. 3xFLAG-PilZ is fully functional. Natural transformation assays of indicated strains. Statistical significance was determined using log-transformed transformation frequencies. Each data point represents a biological replicate (n = 3), and bar graphs indicate the mean ± SD. Statistical significance was determined using Dunnett's multiple comparisons test against the parent strain for each data set. ****$p < 0.0001$; *$p < 0.05$

### Supplemental Figure S4

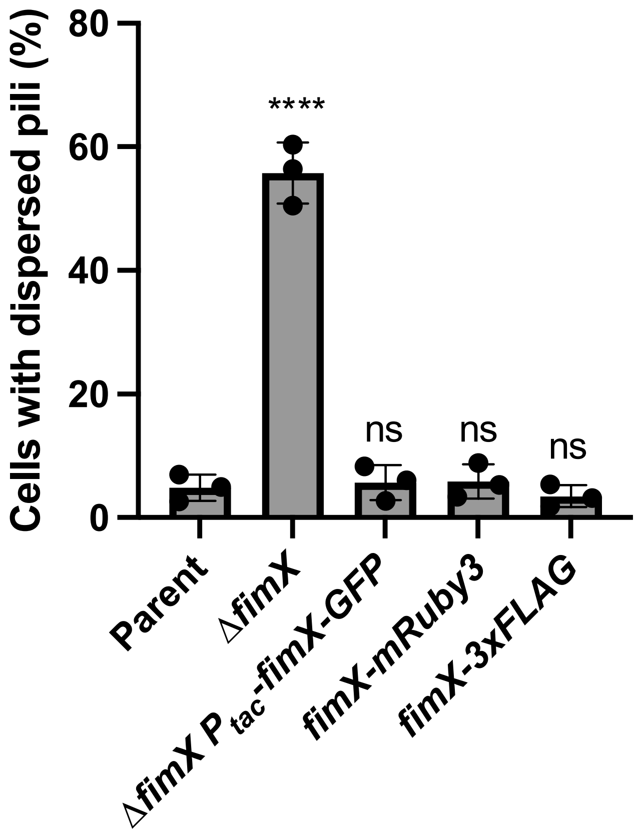


Supplemental Figure S4. All FimX C-terminal fusion strains used in this study are fully functional. Percent of cells with mislocalized or dispersed T4P in indicated strains in a Δ*pilT* background. Each data point represents a biological replicate (n = 3), and bar graphs indicate the mean ± SD. A minimum of 50 cells were analyzed per replicate. Statistical significance was determined using Dunnett's multiple comparisons test against the parent strain for each data set. ****$p < 0.0001$. ns, not significant.

### Supplemental Figure S5

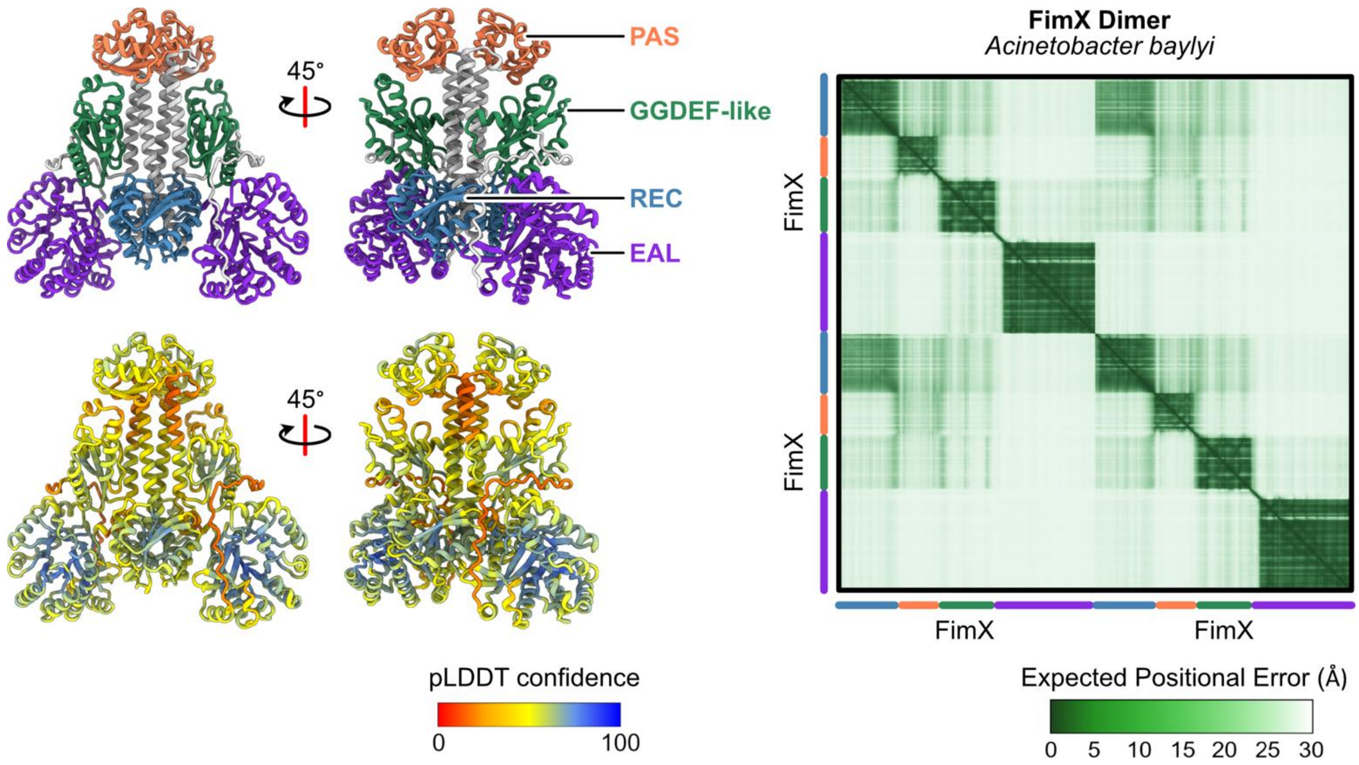


Supplementary Figure S5. AlphaFold3 prediction confidence of the *A. baylyi* FimX dimer. (left) Highest-ranking AlphaFold3 predicted model of *A. baylyi* FimX depicted in two side views and colored by domain organization as in Figure 4A and predicted local distance difference test (pLDDT) scores, ranging from 0-100. (right) Residue-residue predicted alignment error (PAE) plot. The values range from 0-30 Å. Corresponding FimX domains associated with the plot is shown.

### Supplemental Figure S6

```
ADP1fimX      MRNGLLAKKIKRSEIRVLFIDDNQLRYNQVIDLLSAKDYNVHAVLLDDLVSFEKQLKQSW      60
PAO1fimX      -------MAIEKKTIRLLILEDSQNEAERLVSLFRNAGHATRVHRLTSPEDLAETLQQSW      53
                     *::. **:*:::*.* . ::::.*:   .: .:.  * .  .: : *:***

ADP1fimX      DIIIFGRAYDLELTHTISLVHQTPTRQLPILLLDFEQLSPAIQIDFVQKGIYDFLDLTHH      120
PAO1fimX      DLLIAAPTSECLEPGEALATLRRQSRDIPFIQLVADNSSDAIT-DALMLGAQDALPQGED      112
              *::* . : :          . :  :*::*:: *  :: * **  * :  *  * *   ..

ADP1fimX      EAFYIIIARAYTYSKLLQTEQQLNHALKEAHSRTQSLVKQSRKAVAIIEEGIHIQANAEY      180
PAO1fimX      ERLVLVANRELAGLAERRARRAAELALREAEKRCQLLLDSSVDAITYVHDGMHIYANRSY      172
              * : ::  *  :     ::.:  : **:**..* * *:..* .*:: :.:*:** ** .*

ADP1fimX      LALFQLQDQDDLIGIPLLDILQPENVAEFKQAFKKISLGHLEKNRLDMISKNPAVAS---      237
PAO1fimX      MALFGYEDAEELEGLPVIDLIASCDQGAFKDFLKGYQNDQRQT---ELVCTGVKLDGQEF      229
              :***  :* ::* *:*::*::   : . **: :*  . .: :.   :::...  : .

ADP1fimX      SNPLKINFLPHDDAETIQLTIDCDVEQLNLPVLQLE----DST---LEKQLETDLVDSVN      290
PAO1fimX      KARISLSAATYDGEPCIQVVIRGEVDNAELEEKLREVSSQDPVTGLYNRSHFLDLMDAAV      289
              .  :.:.   :*.   **:.*  :*:: :*     *    * .    ::.   **:*:.

ADP1fimX      QHLKITQAKLNSLVLFMPLFNYQQLFNY-DWHQIKQFFSELET-ELNAFQSDCPLRKLEN      348
PAO1fimX      QQA-V-TARKPSTLAYIHLNGYPSLQADHGLSGIDLLLGQLAGLMREQFGEEADLARFGD      347
              *:  :  *:  * : :: * .* .*    .   *. ::.:*     : * .:. * :: :

ADP1fimX      GSYLTIFQASSIEALRTQLMQLGQRISEKFPCSLHELDF---KVELKSGYSLIPSMIESE      405
PAO1fimX      SIFAALFKGKTPEQAQAALQRLL----KKVENHLFELNGRSAQATLSIGVAGLD---EKT      400
              . : ::*:..: *  :: * :*     :*.   *.**:    :. *. * : :    *.

ADP1fimX      ESLENYLNKA-----------LHQVLPYKDKQPVQSFAIPNVKLKTEQKPIEPFTAMADL      454
PAO1fimX      AKAQDVMNRAHRCADDAARKGGSQIKQYN---PAEELAAA-----------AQRGDVIAI      446
               . :: :*:*             *:  *:   *.:.:*                   :  :

ADP1fimX      IKQKFNQGNIVLKYQQLYDKQDQNLYIYEVTRGYY-------DQEHWHCLDDQAGLDNHI      507
PAO1fimX      LQQALETNSFRLLFQPVISLRGDSHENYEVLLRLLNPQGQEVPPAEFLHAAKEAGLAEKI      506
              ::* :: ..: * :* : . :.:.   ***                .:    .:*** ::*

ADP1fimX      DLILQIDRFTLETATKQLKQFINQYPSAKLIINLSEHILKLPQLDKLIEQLLNILRSQEK      567
PAO1fimX      D------RWVILNSIKLLAEHRAKGHQTKLFVHLSSASLQDPGLLPWLGVALKAARLP-P      559
              *      *:.: .: * * :.  :  .:**::**.  *: * *   :   *:  *

ADP1fimX      YPVILQFSEAAFMQNLANAQQQIRQLHQKGVLISVRDFGVNASSPVFLHQIEVDYLQLHH      627
PAO1fimX      ESLVFQISEADATSYLKQAKQLTQGLATLHCQAAISQFGCSLNPFNALKHLTVQFIKIDG      619
                :::*:***   . * :*:*  : *        :: :** . .    *::: *:::::.

ADP1fimX      QLSALLHQENQLQELQQKIDSFKASKSVEMILPELNDVNIFANAWNVSTRYLKGSYFQGK      687
PAO1fimX      SFVQDLNQVENQEILKGLIAELH-EQQKLSIVPFVESASVLATLWQAGATYIQGYYLQGP      678
              .:   *:* :: : *:  * .:: .:.   *:* ::...::*. *:..: *::* *:**

ADP1fimX      LDRLVDV---QDQ*    697
PAO1fimX      SQAMDYDFSSGDE*    691
               : :       *:*
```

Supplementary Figure S6. Pairwise sequence alignment between *A. baylyi* (ADP1) and *P. aeruginosa* (PAO1) FimX. Green boxes show the degenerate “GGDEF” motif while purple boxes depict residues involved in cdG binding in *P. aeruginosa* FimX that diverged in *A. baylyi*.

### Supplemental Figure S7

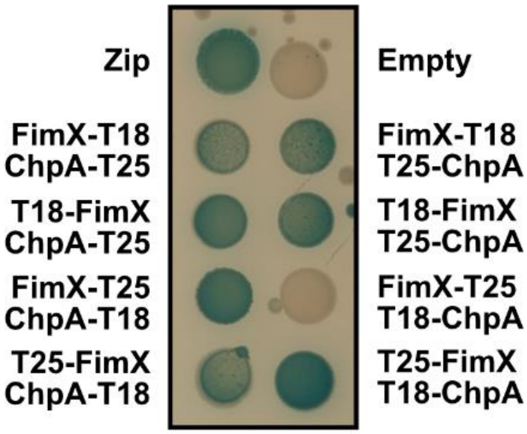


Supplementary Figure S7. FimX directly interacts with ChpA. All combinations of BACTH interactions between FimX and ChpA fused to the N- or C-terminus of the T25 and T18 domains of adenylate cyclase following incubation at 30 °C for 48 hours and subsequently at 4 °C for 24 hours on LB agar plates containing X-Gal. Leucine zipper motif (zip) fused to T18 and T25 was used as positive control while empty vectors of T18 and T25 alone (empty) was the negative control.
